# Determinants of accelerated metabolomic and epigenetic ageing in a UK cohort

**DOI:** 10.1101/411603

**Authors:** Oliver Robinson, Marc Chadeau Hyam, Ibrahim Karaman, Rui Climaco Pinto, Giovanni Fiorito, He Gao, Andy Heard, Marjo-Riitta Jarvelin, Mathew Lewis, Raha Pazoki, Silvia Polidoro, Ioanna Tzoulaki, Matthias Wielscher, Paul Elliott, Paolo Vineis

## Abstract

Markers of biological ageing have potential utility in primary care and public health. We developed an elastic net regression model of age based on untargeted metabolic profiling across multiple platforms, including nuclear magnetic resonance spectroscopy and liquid chromatography-mass spectrometry in urine and serum (almost 100,000 features assayed), within a large sample (N=2,239) from the UK occupational Airwave cohort. We investigated the determinants of accelerated ageing, including genetic, lifestyle and psychological risk factors for premature mortality. The metabolomic age model was well correlated with chronological age (r=0.85 in independent test set). Increased metabolomic age acceleration (mAA) was associated (p<0.0025) with overweight/obesity and depression and nominally associated (p<0.05) with high alcohol use and low income. DNA methylation age acceleration (N=1,102) was nominally associated (p<0.05) with high alcohol use, anxiety and post-traumatic stress disorder, but not correlated with mAA. Biological age acceleration may present an important mechanism linking psycho-social stress to age-related disease.

## Introduction

Ageing can be defined as the “time-dependent decline of functional capacity and stress resistance, associated with increased risk of morbidity and mortality” (Burkle et al., 2015). Environmental stressors, including social adversity (Fiorito et al., 2017; Stringhini et al., 2017), psychological disorders (Chiu et al., 2018; Wolf & Morrison, 2017), and genetic factors(McDaid et al., 2017) may influence the ageing process, leading to differing ageing rates. Traditionally, quantitative assessment of “the rate of ageing” relies on the analysis of mortality curves of populations. However, at the level of a living individual, this method does not allow assessment of the state of ageing (i.e. the state of the functional decline) and a prediction of the risk of morbidity and remaining life expectancy. Therefore, markers of ‘biological age’ (the ageing state typical of one’s chronological age) that can be assessed at any point in the lifespan therefore, may have enormous potential in both personalised medicine and public health. Since ageing is a process that affects almost all tissues and organs of the body and involves cross-talk between multiple physiological systems, there has been increased research into composite markers of ageing, involving multiple parameters (Jylhävä, Pedersen, & Hägg, 2017). Levine (Levine, 2013) employed 10 biomarkers representing multiple systems to develop a biological age score, that could better predict mortality than chronological age. Belsky *et al*. (Belsky et al., 2015) used a similar selection of biomarkers measured longitudinally in young adults to develop a biological age score and found that increased pace of ageing was associated with measures of functional decline such as cognitive ability. Modern ‘omics’ platforms have provided new opportunities for the systematic assessment of biological ageing. For example, Horvath (Horvath, 2013) and Hannum *et al*. (Hannum et al., 2013) employed genome-wide DNA methylation to develop highly predictive models of age based on multiple methylated CpG loci.

Furthermore, it has been shown that ‘age acceleration’, defined as having a greater DNA methylation age than chronological age, is associated with multiple risk factors of mortality such as low social class, smoking, and alcohol use (Fiorito et al., 2017) and is predictive of mortality (B. H. Chen et al., 2016; Dugue et al., 2018). Agnostic metabolomics is a promising candidate technology to develop biomarkers of ageing. Several metabolomic studies have found strong associations between numerous metabolites and age, although in a limited sample size (Chaleckis, Murakami, Takada, Kondoh, & Yanagida, 2016; Rist et al., 2017) or through employing targeted analyses that give limited coverage of the full metabolome (Auro et al., 2014) (Hertel et al., 2016; Yu et al., 2012). Only the study of Hertel *et al*. (Hertel et al., 2016) combined a small set of markers to provide an overall assessment of biological ageing, observing that the predicted metabolomic age was associated with time to death, after adjustment for chronological age and other risk factors.

In the present study, we have employed untargeted metabolomics across multiple analytical platforms, providing unprecedented metabolome coverage (almost 100,000 features assayed), to develop a predictive model of age, within a large sample from the UK occupational Airwave cohort. A second cohort was used for longitudinal validation of selected metabolic age predictors. We have explored the relationship between metabolomic age and DNA methylation age and lifespan associated genetic factors. Furthermore, we have investigated the determinants of accelerated ageing, focussing on risk factors of premature mortality, including the WHO “25 x 25” risk factors (World Health Organisation, 2013) (hypertension, diabetes, obesity, smoking, alcohol use and physical inactivity) and socio-economic and psychological risk factors (income, depression, anxiety, post-traumatic stress disorder (PTSD)). We show that obesity and depression are associated with accelerated metabolomic ageing.

## Results

### Building and validation of the metabolomic age model

The study population included 2,238 participants of the AIRWAVE cohort that had full metabolomic data. 60.5% of participants were male and mean age was 41.24 years (SD: 9.1, range: 19.2 – 65.2 years). Most participants (97.5%) were of white British ethnicity and 27.8% of participants were educated to degree level. The demographic characteristics of this sample are representative of the wider cohort(Elliott et al., 2014). Further covariate information is provided in table 1.

**Table 1:** Demographic and covariate information of participants with metabolomic data, and bivariate associations with metabolomic age acceleration.

|  |  | BIVARIATE ASSOCIATION<br>WITH METABOLOMIC AGE<br>ACCELERATION |  |  |
| --- | --- | --- | --- | --- |
| | | N (%) or Mean (SD) | $\beta$ (95% CI) | p value |
| **DEMOGRAPHIC AND NCD RISK FACTORS** |  |  |  |  |
| **AGE** | years | 41.24 (9.1) | - | - |
| **SEX** | Female | 884 (39.5) | - | - |
|  | Male | 1354 (60.5) | -0.01 (-0.26, 0.24) | 0.94 |
| **MARITAL STATUS** | Married/cohabiting | 1751 (80.2) | - | - |
|  | other | 431 (19.8) | 0.11 (-0.2, 0.41) | 0.5 |
| **ETHNICITY** | White | 2181 (97.5) | - | - |
|  | other | 56 (2.5) | -0.37 (-1.14, 0.4) | 0.35 |
| **BODY MASS INDEX** | <25 | 731 (32.7) | - | - |
|  | >=25 & < 30 (overweight) | 1041 (46.5) | 0.5 (0.23, 0.77) | 0.00035 |
|  | >=30 (obese) | 466 (20.8) | 1.01 (0.67, 1.34) | 4.6E-09 |
| **DIABETIC STATUS** | Normal | 2157 (96.4) | - | - |
|  | Diabetic | 80 (3.6) | 0.78 (0.13, 1.42) | 0.019 |
| **HYPERTENSION** | No | 1540 (68.8) | - | - |
|  | Yes | 697 (31.2) | 0.20 (-0.06, 0.46) | 0.13 |
| **INCOME** | High | 879 (39.4) | - | - |
|  | Medium | 812 (36.4) | 0.26 (-0.01, 0.54) | 0.062 |
|  | Low | 538 (24.1) | 0.2 (-0.11, 0.51) | 0.2 |
| **ALCOHOL CONSUMPTION** | None | 174 (7.8) | - | - |
|  | Moderate | 1876 (83.9) | 0.13 (-0.32, 0.58) | 0.57 |
|  | Heavy | 187 (8.4) | 1.02 (0.42, 1.62) | 0.00085 |
| **SMOKING** | Non-smoker | 1539 (68.8) | - | - |
|  | Former smoker | 477 (21.3) | 0.41 (0.11, 0.7) | 0.0077 |
|  | Current smoker | 221 (9.9) | -0.23 (-0.64, 0.18) | 0.28 |
| **PHYSICAL ACTIVITY** | High | 1305 (58.3) | - | - |
|  | Moderate | 585 (26.1) | -0.21 (-0.5, 0.07) | 0.14 |
|  | Low | 348 (15.5) | 0.21 (-0.13, 0.56) | 0.23 |
| **PSYCHOLOGICAL FACTORS** |  |  |  |  |
| **DEPRESSION DIAGNOSIS** | Normal | 1545 (69.1) | - | - |
|  | Minimal symptoms | 501 (22.4) | 0.45 (0.16, 0.74) | 0.0023 |
|  | Depression | 191 (8.5) | 0.91 (0.47, 1.34) | 4.5E-05 |
| **ANXIETY DIAGNOSIS** | Normal | 1733 (79) | - | - |
|  | Borderline | 274 (12.5) | 0.20 (-0.17, 0.56) | 0.3 |
|  | Anxiety case | 188 (8.6) | 0.52 (0.09, 0.96) | 0.019 |
| **EXPERIENCED TRAUMA IN LAST 6 MONTHS** | No | 1896 (84.7) | - | - |
|  | Yes, without PTSD | 261 (11.7) | 0.14 (-0.24, 0.51) | 0.47 |
|  | Yes, with PTSD | 81 (3.6) | 0.42 (-0.22, 1.07) | 0.2 |
| **DIETARY VARIABLES** |  |  |  |  |
| **FISH CONSUMPTION** | None | 541 (28.6) | - | - |
|  | Medium | 697 (36.8) | 0.25 (-0.08, 0.58) | 0.14 |
|  | High | 654 (34.6) | 0.51 (0.17, 0.84) | 0.0029 |
| **FRUIT CONSUMPTION** | Low | 634 (33.5) | - | - |
|  | Medium | 637 (33.7) | 0.11 (-0.22, 0.43) | 0.51 |
|  | High | 621 (32.8) | 0.11 (-0.21, 0.44) | 0.50 |
| **RED MEAT CONSUMPTION** | Low | 580 (30.7) | - | - |
|  | Medium | 704 (37.2) | 0.21 (-0.11, 0.54) | 0.20 |
|  | High | 608 (32.1) | 0.12 (-0.22, 0.45) | 0.49 |
| **VEGETABLE CONSUMPTION** | Low | 23 (1.2) | - | - |
|  | Medium | 627 (33.1) | 0.17 (-0.16, 0.5) | 0.31 |
|  | High | 617 (32.6) | 0.08 (-0.24, 0.41) | 0.61 |
| **WHOLE GRAIN CONSUMPTION** | Low | 648 (34.2) | - | - |
|  | Medium | 604 (31.9) | 0.23 (-0.1, 0.56) | 0.17 |
|  | High | 661 (34.9) | 0.17 (-0.15, 0.49) | 0.3 |
| **DASH SCORE** | <20 (least healthy) | 344 (18.2) | - | - |
|  | ≥20 and <23 | 357 (18.9) | 0.54 (0.1, 0.97) | 0.015 |
|  | ≥23 and <25 | 297 (15.7) | 0.18 (-0.28, 0.63) | 0.45 |
|  | ≥25 and <28 | 396 (20.9) | 0.67 (0.24, 1.09) | 0.002 |
|  | > 28 (most healthy) | 498 (26.3) | 0.23 (-0.17, 0.64) | 0.26 |
| **MEDITERRANEAN DIET SCORE** | continuous (1-10) | 4.73 (1.82) | -0.03 (-0.11, 0.04) | 0.35 |
| **CLINICAL BIOMARKERS** |  |  |  |  |
| **SYSTOLIC BLOOD PRESSURE** | mmHg | 130.85 (15.21) | 0.01 (0, 0.01) | 0.082 |
| **DIASTOLIC BLOOD PRESSURE** | mmHg | 79.67 (10.1) | 0.01 (-0.01, 0.02) | 0.28 |
| **PULSE** | beats/minute | 70.37 (11.44) | 0 (-0.01, 0.01) | 0.76 |
| **FIBRINOGEN** | g/L | 3.87 (0.88) | 0.02 (-0.12, 0.16) | 0.75 |
| **PROTHROMBIN TIME** | seconds | 13.64 (1.41) | -0.08 (-0.17, 0) | 0.058 |
| **C-REACTIVE PROTEIN** | mg/l | 1.92 (3.1) | 0.02 (-0.02, 0.06) | 0.40 |
| **CREATININE** | μmol / L | 92.35 (12.73) | 0.01 (0, 0.02) | 0.017 |
| **TOTAL CHOLESTEROL** | mmol/l | 5.25 (1.01) | 0.41 (0.29, 0.53) | 9.6E-12 |
| **HIGH DENSITY LIPOPROTEIN** | mmol/L | 1.5 (0.39) | 0.06 (-0.24, 0.37) | 0.68 |
| **Γ-GLUTAMYL TRANSFERASE** | U/L | 31.35 (24.06) | 0.01 (0.01, 0.02) | 1.7E-05 |
| **APOLIPOPROTEIN A** | g/L | 1.35 (0.3) | -0.14 (-0.54, 0.25) | 0.48 |
| **APOLIPOPROTEIN B** | g/L | 0.93 (0.23) | 1.55 (1.04, 2.07) | 3.9E-09 |
| **% GLYCATED HAEMOGLOBIN** | % | 5.63 (0.48) | 0.32 (0.07, 0.57) | 0.013 |
| **UREA** | μmol / L | 5.04 (1.17) | 0.08 (-0.02, 0.18) | 0.12 |

Metabolomic data were acquired from both urine and serum samples using multiple Nuclear Magnetic Resonance Spectroscopy (NMR) and Ultra-Performance Liquid Chromatography - Mass Spectrometry (UPLC-MS) platforms, providing in total nine different metabolomic data types (table s1). For purposes of constructing the main predictive model of age through elastic net regression, these data types were combined into one metabolomic dataset, giving a total of 98,824 metabolic features.

In the first stage of model building, an analysis by metabolomic platform (sequentially leaving on platform out each time) indicated that predictive performance (minimisation of mean squared error in 10-fold cross validation) was improved through using only the four following platforms (figure s1): Bruker IVDr Lipoprotein Subclass Analysis derived from NMR in serum (“sBiLISA”), lipid-targeted reverse-phase UPLC-MS in positive mode in serum (“sLPOS”), reverse-phase UPLC-MS in positive mode in urine (“uRPOS”) and hydrophilic interaction UPLC-MS in positive mode in urine (“uHPOS”) to give a total of 28941 metabolic features. The final predictive model selected 525 predictors from across this set (see table s2 for list of predictors along with table s3 annotation information), including 8 lipoprotein subclasses from sBiLISA and 219, 104 and 194 features (retention time-m/z pairs) from the sLPOS, uHPOS and uRPOS platforms respectively. The model predicted age with high accuracy (mean absolute error, (MAE) = 1.47 years) in the building data set (80% of data n= 1790), with a correlation between chronological age and predicted age of 0.96 (figure 1a). When this model was applied to the independent validation dataset, consisting of the remaining 20% of study participants (N = 448), the MAE was 3.80 years and the correlation between predicted age and chronological age was 0.85 (figure 1a).

**Figure 1:**
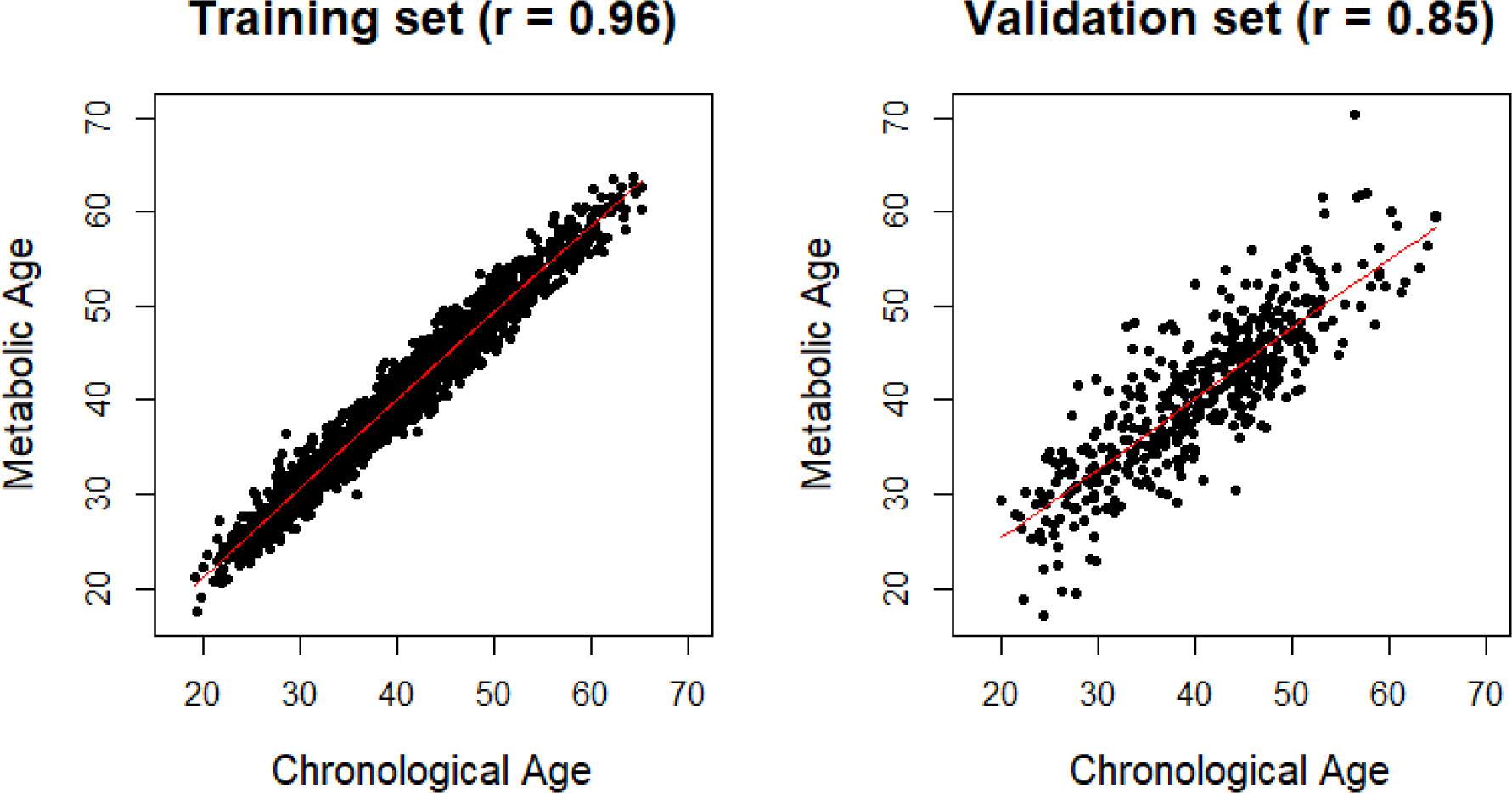
Summary of metabolomic age prediction model. A: Predicted age plotted against chronological age in training set. Pearson’s correlation coefficient (r) is shown. B: Predicted age plotted against chronological age in test set. Pearson’s correlation coefficient (r) is shown

Pathway enrichment analysis, using the *Mummichog* algorithm performed across the UPLC-MS derived model predictors, identified enrichment (p < 0.05) in eleven metabolic pathways (table 2): Vitamin E metabolism, Tryptophan metabolism, CoA Catabolism, Urea cycle/amino group metabolism, Lysine metabolism, Carnitine shuttle, Vitamin B5 - CoA biosynthesis from pantothenate, Biopterin metabolism, Drug metabolism - cytochrome P450, Tyrosine metabolism, and Aspartate and asparagine metabolism.

**Table 2:**
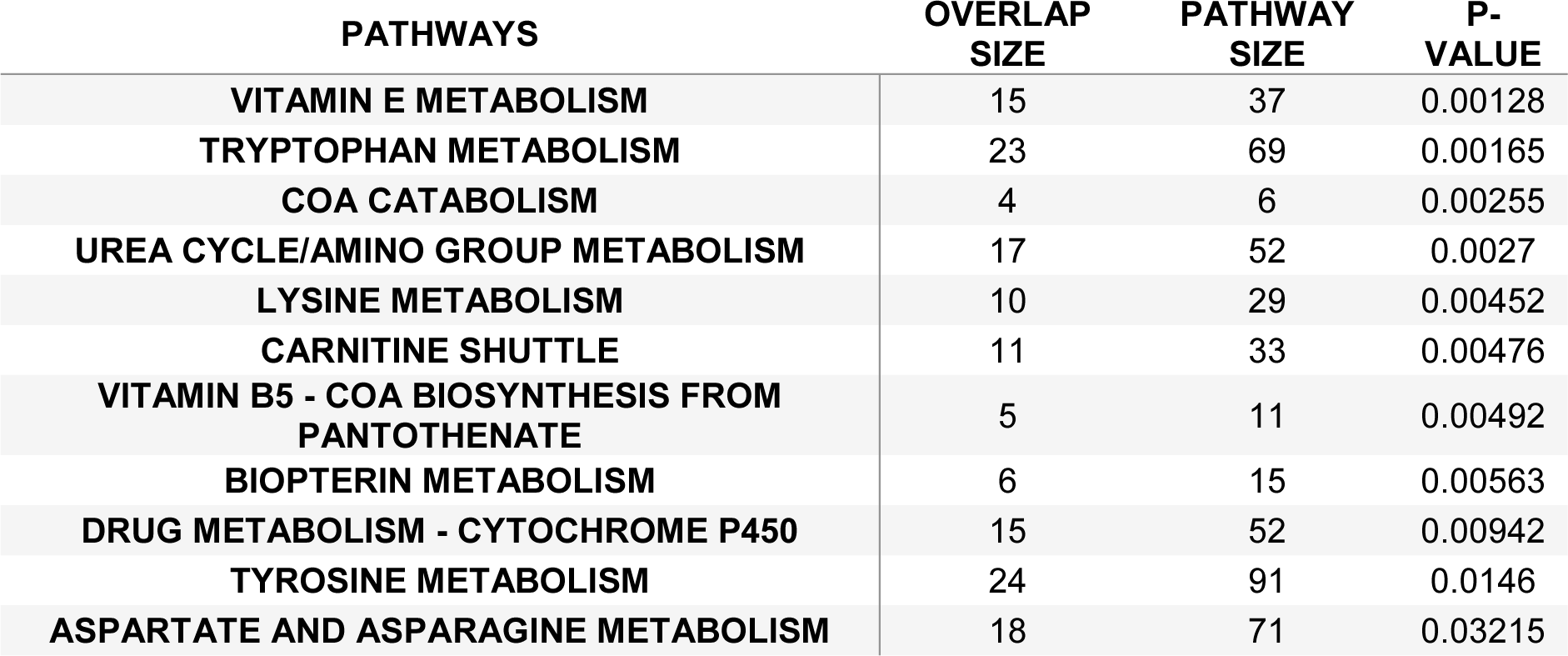
Significantly enriched metabolic pathways among metabolomic age predictors.

| PATHWAYS | OVERLAP<br>SIZE | PATHWAY<br>SIZE | P-VALUE |
| --- | --- | --- | --- |
| VITAMIN E METABOLISM | 15 | 37 | 0.00128 |
| TRYPTOPHAN METABOLISM | 23 | 69 | 0.00165 |
| COA CATABOLISM | 4 | 6 | 0.00255 |
| UREA CYCLE/AMINO GROUP METABOLISM | 17 | 52 | 0.0027 |
| LYSINE METABOLISM | 10 | 29 | 0.00452 |
| CARNITINE SHUTTLE | 11 | 33 | 0.00476 |
| VITAMIN B5 - COA BIOSYNTHESIS FROM<br>PANTOTHENATE | 5 | 11 | 0.00492 |
| BIOPTERIN METABOLISM | 6 | 15 | 0.00563 |
| DRUG METABOLISM - CYTOCHROME P450 | 15 | 52 | 0.00942 |
| TYROSINE METABOLISM | 24 | 91 | 0.0146 |
| ASPARTATE AND ASPARAGINE METABOLISM | 18 | 71 | 0.03215 |

We examined concentration changes of nine metabolites included in our age prediction model, that were available in an independent cohort, the Northern Finnish Birth Cohort 1966, that had serum NMR metabolomic data measured at two ages, 31 and 46 yrs, among 2144 individuals. Seven of these metabolites (77%) changed significantly with age (p < 0.05), in the same direction as predicted in the metabolomic age model (table 3).

**Table 3:**
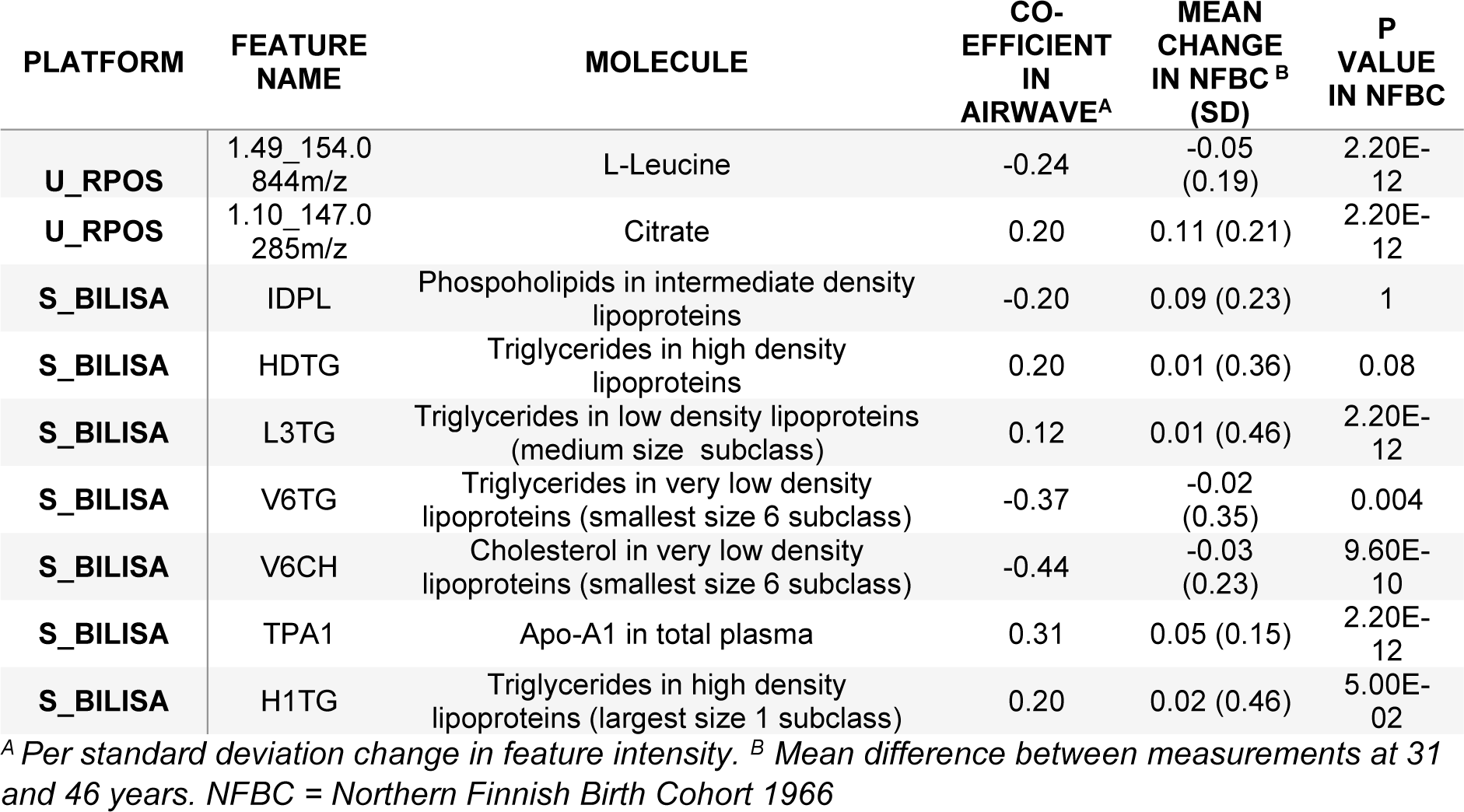
Validation of selected metabolomic age predictors in the Northern Finnish Birth Cohort.

| PLATFORM | FEATURE<br>NAME | MOLECULE | CO-<br>EFFICIENT<br>IN<br>AIRWAVE<sup>A</sup> | MEAN<br>CHANGE<br>IN NFBC<sup>B</sup><br>(SD) | P<br>VALUE<br>IN NFBC |
| --- | --- | --- | --- | --- | --- |
| U_RPOS | 1.49_154.0<br>844m/z | L-Leucine | -0.24 | -0.05<br>(0.19) | 2.20E-12 |
| U_RPOS | 1.10_147.0<br>285m/z | Citrate | 0.20 | 0.11 (0.21) | 2.20E-12 |
| S_BILISA | IDPL | Phospholipids in intermediate density lipoproteins | -0.20 | 0.09 (0.23) | 1 |
| S_BILISA | HDTG | Triglycerides in high density lipoproteins | 0.20 | 0.01 (0.36) | 0.08 |
| S_BILISA | L3TG | Triglycerides in low density lipoproteins (medium size subclass) | 0.12 | 0.01 (0.46) | 2.20E-12 |
| S_BILISA | V6TG | Triglycerides in very low density lipoproteins (smallest size 6 subclass) | -0.37 | -0.02<br>(0.35) | 0.004 |
| S_BILISA | V6CH | Cholesterol in very low density lipoproteins (smallest size 6 subclass) | -0.44 | -0.03<br>(0.23) | 9.60E-10 |
| S_BILISA | TPA1 | Apo-A1 in total plasma | 0.31 | 0.05 (0.15) | 2.20E-12 |
| S_BILISA | H1TG | Triglycerides in high density lipoproteins (largest size 1 subclass) | 0.20 | 0.02 (0.46) | 5.00E-02 |

<sup>A</sup> Per standard deviation change in feature intensity. <sup>B</sup> Mean difference between measurements at 31 and 46 years. NFBC = Northern Finnish Birth Cohort 1966

### Metabolomic age, DNA methylation and genetic predictors of longevity

DNA methylation age was assessed for 1102 participants. Demographic characteristics for this sample were similar to those for participants with metabolomic age available (table s4). DNA methylation age predicted chronological age with a MAE of 4.37 years. DNA methylation age was strongly correlated with chronological age (r=0.91, figure 2a) and metabolomic age (n = 837, r = 0.85, figure 2b). Age acceleration scores were derived for both DNA methylation age acceleration (DNAmAA) and metabolomic age acceleration (mAA), as the difference, at a given age, between actual and predicted age. However, no correlation was observed between DNAmAA and mAA (r = 0.02, figure 2c).

**Figure 2:**
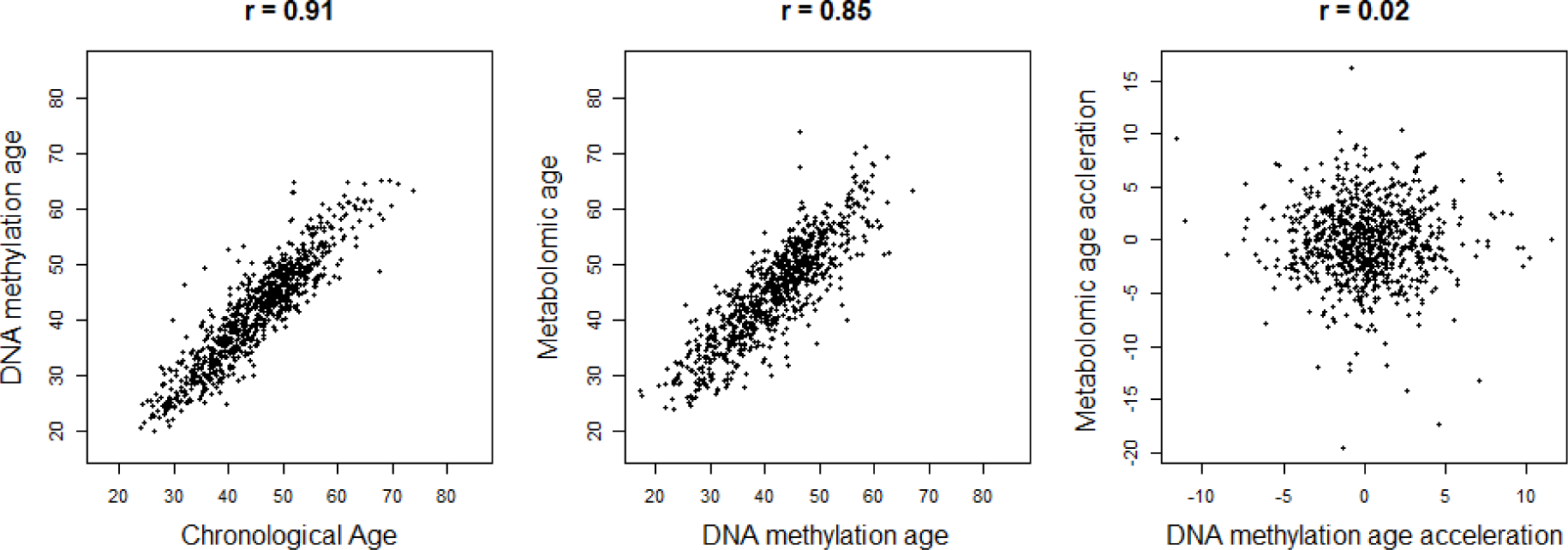
Relationships between different age measures. A. DNA methylation age plotted against chronological age. B: metabolic age plotted against DNA methylation age. C. MetAA plotted against DNAmethAA. Pearson’s correlation coefficients (r) are shown

Table s5 shows mean age acceleration scores by genotype for 10 single nucleotide polymorphisms (SNPs), that have robust associations with parental attained age, identified in two recent studies within the UK Biobank (Joshi et al., 2017; Pilling et al., 2017).

Associations of borderline nominal significance that were consistent with genetic effects on lifespan (i.e. age acceleration increases with genotype associated with shorter lifespan or *visa versa*) were observed only for a SNP in *APOE* locus with both mAA (p = 0.05) and DNAmAA (p =0.07). A weighted genetic risk score (GRS) for shortened lifespan (derived from these 10 SNPs) increased mAA by 1.55 (95% confidence interval (CI): -1.71, 4.81) and DNAmAA by 2.87 (95% CI: -1.56, 7.29) per GIS unit, although this was not statistically significant (p=0.35 with mAA and p=0.20 with DNAmAA).

### Risk factors of age acceleration

In bivariate analyses (table 1) we observed increased mAA (p < 0.05) among participants who were diabetic, heavy drinkers, overweight or obese, former smokers, or were suffering from depression, anxiety or PTSD. Clinical biomarkers associated with mAA included creatinine, total cholesterol, γ-glutamyl transferase (GGT) apolipoprotein B and glycated haemoglobin (%HBa1C). Regarding dietary intake in the week prior to sampling, those who reported high fish consumption and those in the second or fourth quintiles of the DASH score (compared to the first quintile, the least healthy dietary pattern) also had increased mAA. In bivariate analyses with DNAmAA (table s4), sex was associated at p<0.05, with an increase in DNAmAA of 0.89 (interpretable as years of increase in DNA methylation age, 95% CI: 0.47, 1.30) in men compared to women. Clinical biomarkers associated with DNAmAA in bivariate analyses included creatinine, high density lipoproteins, GGT and apolipoprotein A.

Table 5 shows adjusted associations with mAA and DNAmAA for non-communicable disease and psychological risk factors (adjusted for sex, ethnicity, study centre, income, hypertension, diabetes, BMI, smoking, alcohol intake, physical activity, DASH score and fish consumption). We observed nominally significant increases (p<0.05) in mAA with overweight, obesity, heavy drinking, lower income, depressive symptoms and depression, ranging from 0.35 (95% CI: 0.01, 0.69) for low income compared to those with high income, to 0.97 (95% CI: 0.57, 1.37) for obesity compared to those of normal weight. Significant increases in DNAmAA were observed with heavy drinking, anxiety and PTSD, ranging 0.92 (95% CI: 0.03, 1.80) for anxiety compared to those without anxiety symptoms, to 2.15 (95% CI: 0.31, 4.00) for PTSD compared to those who had not experienced trauma in the past six months. Associations between overweight/obesity and depression and mAA remained significant (p < 0.0025) after correction for multiple testing.

**Table 5:**
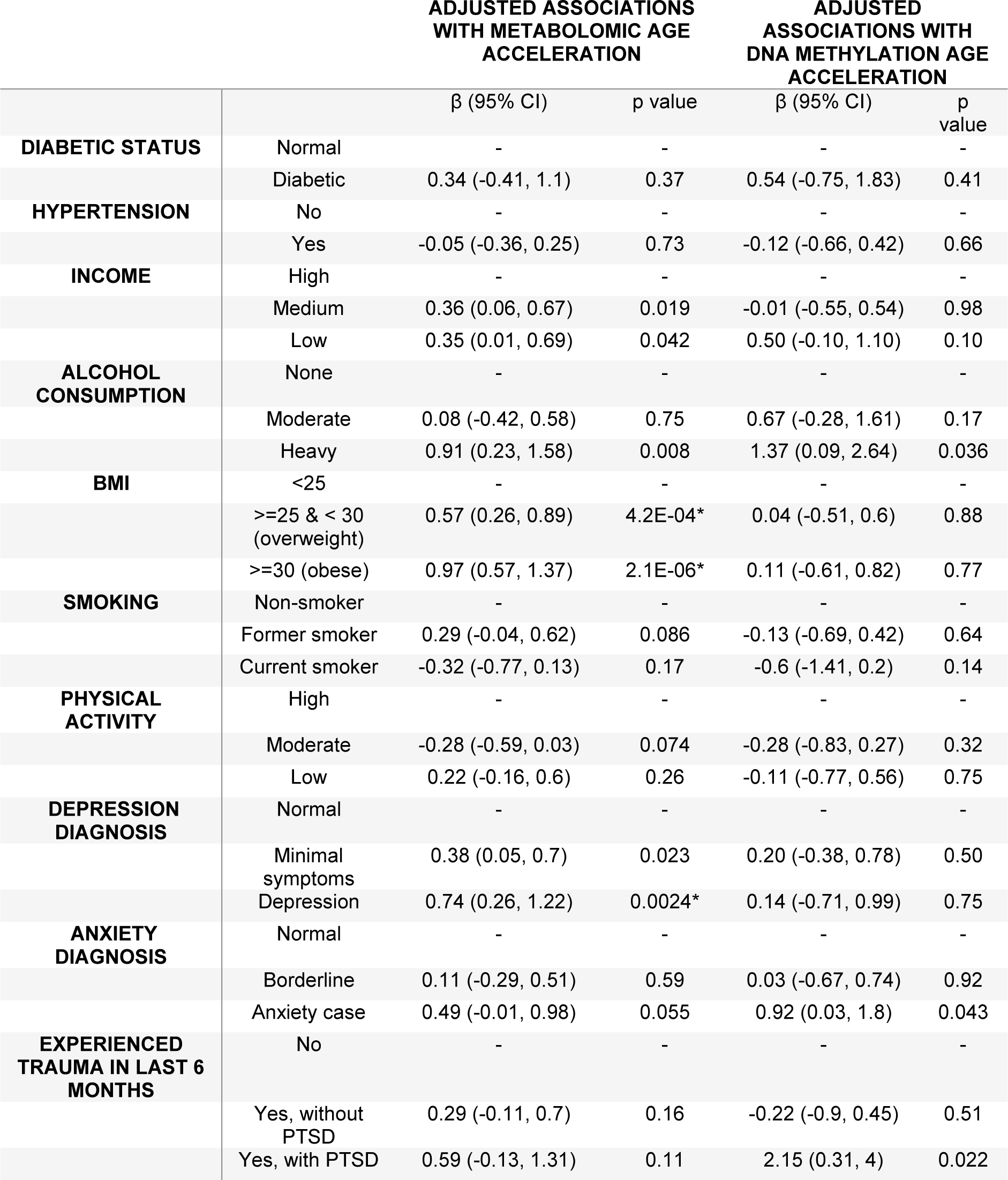
Adjusted associations between disease risk factors and age acceleration scores.

Models adjusted for sex, ethnicity, study centre, income, hypertension, diabetes, BMI, smoking, alcohol intake, physical activity, DASH score and fish consumption. **\*** indicates p values that pass multiple testing correction.

## Discussion

In an important proof of principle study, we have demonstrated in a large nationwide cohort study that metabolomic profiling may be used to predict chronological age with high accuracy among working age adults. We employed a wide range of metabolomic platforms to provide the broadest metabolome coverage yet presented in population-based studies. We found that metabolomic age acceleration, defined as having a greater predicted metabolomic age than chronological age, was associated, after multiple testing correction, with overweight/obesity and depression and nominally associated with low income and high alcohol intake. We did not observe an association between epigenetic age acceleration and metabolomic age acceleration, suggesting these measures capture separate aspects of the ageing process. We observed a different pattern of risk factors nominally associated with epigenetic accelerated ageing including being male, heavy drinking, anxiety and PTSD.

The correlation between chronological and predicted age, of our measure of metabolomic ageing (r= 0.86 in the validation dataset), was somewhat lower than that of the Hannum epigenetic age clock in our cohort (r= 0.91) but greater than reported for other biological ageing markers, including the measure based on urinary NMR data (Hertel et al., 2016)(r = 0.53 in men and 0.61 in women in validation dataset), the blood transcriptomic clock (Peters et al., 2015) (r = 0.35- 0.74 depending on cohort) and telomere length (r ∼ 0.3 (Muezzinler, Zaineddin, & Brenner, 2013)). Biological ageing markers aim to better capture the body’s rate of decline or physiological breakdown than chronological age itself and should therefore also be more predictive of mortality and age-related disease. The associations we observed between accelerated metabolomic ageing and factors known to increase risk of mortality, suggest that metabolomic age may capture this physiological decline.

Strong associations with mAA were observed with overweight and obesity. These conditions are forms of metabolic dysregulation and their additional metabolic burden may increase the rate of decline of the metabolic systems of the body. Genetic predisposition to longevity is associated with low levels of abdominal visceral fat (Sala et al., 2015) and many different conditions that prolong lifespan in animal models also improve obesity-related conditions. Furthermore, obesity has been linked to telomere shortening, and drastic measures to combat morbid obesity like bariatric surgery can actually cause a recovery in telomere length (Laimer et al., 2015). Much is now known about the ageing process at the molecular level primarily from experimental work. López-Otín et al.(López-Otín, Blasco, Partridge, Serrano, & Kroemer, 2013) proposed nine ‘hallmarks of ageing’ that may all be expected to have detectable effects on the metabolome and overlap significantly with the effects of metabolic disorders (López-Otín, Galluzzi, Freije, Madeo, & Kroemer, 2016). For instance, the hallmark ‘deregulated nutrient signalling’ refers to pathways that sense and respond to nutrient availability such as “insulin and IGF1 signalling” (IIS) pathway, which is altered among diabetics.

We observed multiple metabolomic pathways enriched among the predictors of our metabolomic clock that reflect fundamental metabolic processes and are closely related to these hallmarks. We observed enrichment of the pathway related to metabolism of Vitamin E, a potent anti-oxidant and anti-inflammatory agent that protects cell membranes from oxidative damage that can induce genome instability (Claycombe & Meydani, 2001). As a primary hallmark, genome instability has far-reaching and complex consequences including altered nutrient sensing, energy metabolism and redox balance (Garinis, van der Horst, Vijg, & H.J. Hoeijmakers, 2008). Many human progerias are disorders accompanied by the hyperactivation of DNA repair machinery dependent on nicotinamide adenine dinucleotide (NAD+). This hyperactivation leads to NAD+ depletion, resulting in inhibition of the NAD+- dependent nutrient sensor sirtuin 1 (SIRT1) (Fang et al., 2014). Levels of NAD+ are also affected by other factors including circadian rhythm disruption, chronic inflammation (Verdin, 2015) and tryptophan metabolism (also enriched among the metabolomic clock predictors). The functional impairment of SIRT1 (Gomes et al., 2013), which limits expression of nuclear genes encoding mitochondrial proteins, leads directly to the hallmark mitochondrial dysfunction. We observed enrichment among the metabolomic clock predictors of the pathways CoA catabolism, Vitamin B5 - CoA biosynthesis from pantothenate and lysine metabolism, which all maintain acetyl-coA levels necessary for mitochondrial reactions, and carnitine shuttle, which is required for the transport of fatty acids for beta-oxidation in the mitochondria. The enrichment of these pathways suggests the importance of the mitochondrial dysfunction hallmark in our metabolomic ageing model. SIRT1 also contributes to regulating the circadian oscillation of acetyl-coA levels (Sahar et al., 2014) which has been linked to the ageing process (Chang & Guarente, 2013) and epigenetic alterations through acetylation (Su, Wellen, & Rabinowitz, 2016). Mitochondrial fitness further has impact on other ageing hallmarks (López-Otín et al., 2016), including genomic instability (dysfunctional mitochondria are major sources of genotoxic ROS), altered intercellular communication (ROS overgeneration is connected to the secretion of inflammatory mediators) and stem cell exhaustion (which are particularly sensitive to ROS (Ito et al., 2006)). The observed enrichment of the urea cycle and aspartate and asparagine metabolism pathways will also result from perturbation to the Krebs and urea cycles following changes in mitochondrial fitness.

The enrichment of the tryptophan, tyrosine and biopterin metabolic pathways appear to relate to the hallmark ‘altered intercellular communication’. Tyrosine is required for signal transduction through incorporation into protein kinases, while tryptophan and biopterin are necessary for synthesis of neurotransmitters including dopamine, norepinephrine, epinepherine, serotonin and melatonin. Alterations to neurotransmitter levels may underlie the associations we observed between mAA with depressive symptoms and depression. Both psychological distress and major depression had similar hazard ratios for mortality in a recent prospective study (Chiu et al., 2018), which would be consistent with the observed increases in mAA for both depressive symptoms and depression. Anxiety was also associated with increased mAA, albeit a smaller increase than observed for depression. While in this cross-sectional study we cannot disentangle the causal direction between depression and mAA, a study of biological ageing among elderly people found that accelerated biological age was associated with depressive symptoms at baseline and was also predictive of depressive symptoms at follow-up (Brown et al., 2017). Consistent evidence demonstrates a bi-directional association between depression and so-called metabolic syndrome, suggesting common pathological roots (Marazziti, Rutigliano, Baroni, Landi, & Dell’Osso, 2014). Proposed pathophysiological commonalities include abnormal activation of the hypothalamic–pituitary–adrenal (HPA) axis and altered levels of circulating leptin and ghrelin, two peripheral hormones that are classically implicated in the homeostatic control of food intake. A large body of research has investigated the concept of ‘allostatic load’ whereby repeated activation of the HPA axis leads to biological ‘wear and tear’ or physiological decline of downstream metabolic, immune and cardiovascular systems (McEwen & Seeman, 1999). Many studies have demonstrated a link between social adversity (Castagne et al., 2018; Dowd, Simanek, & Aiello, 2009) and allostatic load and it is theorised that chronic stress associated with low socio-economic position leads to prolonged activation of the HPA axis. We observed that lower income was nominally associated with increased mAA, which may similarly be considered to capture physiological decline.

We observed increases in DNAmAA associated with anxiety, PTSD and low income that were generally of greater size than for mAA. Meta-analyses have shown that both PTSD (Wolf & Morrison, 2017) and low socio-economic position (Fiorito et al., 2017) to be associated with increases in DNAmAA. We did not observe any evidence for an association between depression and DNAmAA, suggesting the two ageing measures may be sensitive to separate dimensions of mental health. The DNA methylation clock has been shown to perform well as marker of biological age since it is predictive of all-cause mortality, even after adjusting for chronological age and a variety of known risk factors, and is associated with physical measures of ageing such as frailty and cognitive decline (Horvath et al., 2016). However, other biological ageing markers may add value in capturing different aspects of the ageing process. Peters *et al*. (Peters et al., 2015) reported that transcriptomic age was only moderately correlated with DNA methylation age and the different measures were associated with different ageing phenotypes. Similarly, Belsky *et al*. (Belsky et al., 2016) report only weak correlations between telomere length, DNA methylation age, and a composite biomarker-based measure of biological ageing among young adults. While metabolomic and DNA methylation age were correlated in our study, there was no association between mAA and DNAmAA. DNAmAA has been shown to be predictive of cancer related mortality but not CVD (Dugue et al., 2018; Horvath et al., 2016) while the risk factors associated with mAA suggest it may be predictive of cardio-metabolic related disease. Accelerated transcriptomic age was found to be similarly associated with CVD risk factors, although it was not related to mental health (Peters et al., 2015). Further research into biological ageing may consider combining markers at different levels of biological organisation to provide a more complete picture of the ageing process.

There was suggestive evidence for a small association between the *APOE* gene and both mAA and DNAmAA. This gene has the strongest effects on attained age and is the only genetic variant replicated across populations in studies of human longevity (Partridge, Deelen, & Slagboom, 2018). It plays a key role in lipoprotein metabolism and has been associated with multiple are-related disorders including cognitive decline (De Jager et al., 2012) and Alzheimer’s disease (Marioni et al., 2018). The role of the *APOE* gene in these biological age markers, requires further study in larger, independent populations.

This study had some important limitations. The study was cross-sectional, based on a single biological sampling from participants at a wide range of ages. It is therefore difficult to separate processes relating to the ageing itself from cohort effects associated with the different environment of people at different ages. This is particularly a problem for analyses of the metabolome which contains both endogenous metabolites related to physiological processes such as ageing and short-lived exogenous metabolites related to factors such as diet and medication. Indeed, we observed that fish consumption, which is associated with reduced risk of mortality (Zhao et al., 2016), actually increased mAA, likely due to the confounding of our model by cohort effects. We addressed these points in two ways: Firstly, we validated some of the metabolomic age predictors that were available in an independent cohort at two timepoints (15 years apart) in the early adult life of the same individuals. We found that there were highly significant changes in levels of the majority of metabolites we checked, in the same direction as predicted by the metabolomic age model. Secondly, we adjusted associations with mAA for diet that had been assessed through a food diary in the week prior to sampling. Pathways that were enriched in our model were generally related to physiological processes known to be related to ageing, with the possible exception of the drug metabolism pathway. However, medication history was unavailable in this study. Also, we did not have participants at the oldest ages (>65 yrs). It is known that biological ageing becomes more variable within the elderly and further work is required to test the performance of our modelling approach within older populations.

The use of untargeted metabolomics presents both strengths and limitations. Untargeted analyses reduce the potential to apply the full model in separate metabolomic datasets due to differences in retention time and mass accuracy in different runs of spectral acquisition. Furthermore, full laboratory annotation of all predictors was outside the scope of the present study and may not even be possible for some predictors without current database matches. However, the aim of the study was to develop an overall predictive model to assess metabolic ageing rather than identify individual predictors. Indeed, the nature of the variable selection method used means that an equally valid predictive model can be built on different sets of predictors. We used the *Mummichog* pathway analysis tool to extract information at the pathway level, as the algorithm bypasses laboratory annotation based on the assumption that misidentification will apply equally both to the feature set (metabolites included in the age prediction model) and the reference set (metabolites not selected into the model). The tool has been validated in separate datasets that have also undergone full laboratory annotation (Li et al., 2013). We incorporated a range of MS platforms able to detect both lipophilic and hydrophilic molecules at low concentrations and NMR platforms able to detect larger structures such as lipoproteins that would be destroyed during MS acquisition. We also analysed both serum and urine that contain different sets of metabolites – more lipophilic molecules in serum and more polar molecules that are present at higher concentrations in urine. Together, we were able to assay a large portion of the metabolome that would not be possible with current targeted methods.

Other strengths include the incorporation of genomic and DNA methylation data, the wide age range of participants including those in early adult life where ageing interventions may be most effective (Moffitt, Belsky, Danese, Poulton, & Caspi, 2017), and the use of validated psychological instruments. Future work will assess the effects of mAA on functional ageing measures and other health endpoints and assess metabolomic age in longitudinal, repeat samples.

In conclusion, we have developed a predictive indicator of aging based on broad metabolomic analysis among working age adults. We found that while mAA, the difference between metabolomic and chronological age, was not related to DNAmAA, it was associated with mortality risk factors including overweight/obesity, heavy alcohol use and psycho-social factors including depression and lower income. Biological age acceleration may be an important mechanism linking psycho-social stress to age-related disease. Advances in life expectancies have led to an increased prevalence of age-related morbidities. Targeting the process of ageing itself, through changes in living conditions, behaviours or therapeutic interventions, may help more people experience healthy ageing.

## Materials and Methods

### Cohort and covariate information

The Airwave Health Monitoring Study is an occupational cohort of employees of 28 police forces from across Great Britain. Full details of the cohort and methods are available in Elliott et al (Elliott et al., 2014). The study started recruitment in 2006 and now contains 53,280 participants. The study received ethical approval from the National Health Service Multi-Site Research Ethics Committee (MREC/13/NW/0588). At the baseline health screening, participants underwent health examination, self-completed a computer questionnaire and provided urine and blood samples. Blood samples were spun at the health clinic and the biological samples were stored in a Thermoporter (LaminarMedica) and sent overnight from the clinics for next-day analysis of standard clinical chemistry tests or were frozen at -80 °C long term storage. DNA samples and plasma for metabolomic analysis were extracted from blood collected in EDTA tubes.

Important covariates in the analysis were categorised from self-report or clinical data as follows: Ethnicity was defined as ‘white’ or otherwise. Marital status was defined as living with partner or otherwise. Income was defined as low, medium or high, based on terciles of total net household income after adjustment for the number of dependant household members. Alcohol use was classed as non-drinker, moderate drinker (≤ 14 alcohol units/week for women and ≤ 21 alcohol units/week for men) or heavy drinker (> 14 alcohol units/week for women and >21 alcohol units/week for men). Hypertension was defined as ether reported diagnosis or systolic blood pressure ≥ 140 mmHg or diastolic blood pressure ≥ 90 mmHg. Diabetic status was defined as normal (no diagnosis and HbA1c < 6.5%), or diabetic (diagnosis or HbA1c ≥ 6.5%). Physical activity was defined as low, moderate or high based on the scoring protocol of the International Physical Activity Questionnaire (The IPAQ group, 2016).

### Psychological instruments

The Patient Health Questionnaire – 9 depression questionnaire was used to define participants as “normal (i.e. no depression)”, “minimal symptoms of depression” or as a “depression case” (Kroenke, Spitzer, & Williams, 2001). The Hospital Anxiety and Depression Scale questionnaire was used to assess anxiety levels as “normal (i.e. no anxiety)”, “borderline” and “anxiety case” (Zigmond & Snaith, 1983). Participants were asked if they had experienced a work-related traumatic incident in the previous six months. Those who reported a traumatic incident were then asked to complete a brief screening instrument for post-traumatic stress disorder (PTSD) (Brewin et al., 2002). Participants were thus classed into three categories: “not experienced traumatic incident in past 6 months”, “experienced traumatic incident in past 6 months without leading to PTSD”, and “experienced traumatic incident in past 6 months leading to PTSD”,

### Assessment of Diet

Dietary intake was measured using validated 7-day estimated weight food diaries as fully described previously (Gibson et al., 2017). Nutritional intake was calculated using Dietplan6.7 software (Forestfield Software, Horsham, UK) which is based on the McCance and Widdowson’s 6th Edition Composition of Foods UK Nutritional Data set (UKN) by a team of trained coders trained to match food and drink items to the UKN database code and a portion size.

Energy adjusted average consumption of fruit, vegetables, red meat, processed meat, wholegrain and dairy over the week was categorised into tertiles. For fish, consumption was divided into none, medium (below median consumption among consumers) and high (above median consumption). Two overall dietary scores were calculated: The Dietary Approaches to Stop Hypertension (DASH) diet score divided into quintiles (Fung et al., 2008) and the Mediterranean Diet score as a continuous measure (Trichopoulou, Costacou, Bamia, & Trichopoulos, 2003).

### Metabolomic data acquisition

Metabolomic analysis of serum and urine was performed at the National Phenome Centre, based at Imperial College London. Samples were randomly sorted into batches of 80 and thawed to 4°C, centrifuged to remove particulate matter, and the supernatant dispensed across dedicated 96-well plates for each assay. Study-Reference (SR) samples, a pool of all samples for each matrix in the study, and Long-Term Reference (LTR) samples, a pool of samples external to study, were included in each analytical run to allow for quantification and correction of technical variation. Samples are prepared and analysed daily in batches of 80 study samples with the addition of 4 quality controls (2 SR and 2 LTR). Samples were maintained at 4°C during preparation for, and while awaiting, acquisition.

Acquisition of Nuclear Magnetic Resonance Spectroscopy (NMR) profiles (the NOESY experiment in urine and the CPMG experiment in serum) was conducted as described in (Dona et al., 2014). Lipoprotein parameters were generated by the Bruker B.I.-LISA (Bruker IVDr Lipoprotein Subclass Analysis platform, derived from NMR of serum. Spectra were acquired at 600 MHz with Bruker Ascend 600 magnets and Avance III HD consoles configured to the Bruker IVDr specification (Bruker Corporation, Billerica, MA, USA).

Ultra-Performance Liquid Chromatography - Mass Spectrometry (UPLC-MS) acquisitions were conducted in batches of up to 1000 study-samples, interleaved with alternating SR and LTR samples every five injections (16 per 80 samples), each batch was flanked by a serial dilution of the SR sample to assess linearity of response. Multiple analytical experiments were performed to increase metabolomic coverage. Hydrophilic interaction chromatography was performed in both urine and serum as described in (Lewis et al., 2016). Reversed-phase chromatography was performed on urine samples in both positive and negative modes as described in (Lewis et al., 2016). Lipid-targeted reverse-phase chromatography was applied in serum ionised in both positive and negative modes as described in (Sarafian et al., 2014). All UPLC-MS profiling assays were acquired on Waters G2-S ToF mass spectrometers, with Acquity UPLC chromatography systems (Waters Corporation, Milford, MA, USA).

### Metabolomic data processing

NMR spectra were automatically processed in TopSpin 3.2, followed by a suite of in house scripts (Dona et al., 2014). Each spectrum was automatically checked, before all spectra were aligned to a common reference scale. Analytical quality was further assessed manually on four factors: Line width of less than 0.9 Hz, quality of water-suppression, even baseline signal and accurate chemical shift referencing. Urine samples were referenced to an internal spiked standard 3-(trimethylsilyl)-2,2,3,3-tetradeuteropropionic acid (TSP) at 0 ppm. Plasma samples were referenced to the α-anomeric glucose doublet at 5.233 ppm. Spectra were aligned to a common reference scale, running from 10 to -1 ppm, and interpolated onto a common 20,000 point grid. Lipoprotein parameters were validated according to Bruker’s B.I.-LISA protocols.

Chromatograms and mass spectra instrument raw files were imported into Progenesis QI (Waters Corp. Milford, MA, USA) for retention-time alignment and feature detection. Progenesis QI was configured to align retention time to the central LTR sample of the acquisition. Peak detection was configured with a minimum chromatographic peak width of 0.01 minutes, and automatic noise detection set to the minimum threshold of 1. Peaks arising from isotopes and chemical adducts were automatically resolved according to the observed m/z and chromatographic peak shape, and peaks areas integrated. Further processing and filtering of UPLC-MS profiling datasets was conducted with in-house scripts, and used to account for analytical run-order effects and remove noise from each dataset.

Analytical run-order effects were accounted for with an adaption of the method described in (Zelena et al., 2009). A robust LOWESS regression was generated per-feature, based on the SR samples, in run-order, with the window scaled to include 21 SR samples. The smoothed response values for each feature were then interpolated to the intermediate study sample injections using simple linear interpolation.

Finally, the median intensity of each feature in each analytical batch was aligned. Extracted features spuriously arising from analytical noise were removed from the dataset by a pair of approaches, both applied on a per-feature basis. First, a serial dilution of the study reference sample was used to assess the linearity of responses of each feature. Detected features were correlated to their expected intensity in the dilution series, and those features showing a Pearson’s r of less than 0.7 were excluded from further analysis. Second, the relative standard deviation (RSD) of each feature across the study reference samples was calculated, and those features where the RSD exceeded 30%, or the observed biological variance was less than 1.5 times the RSD, were excluded.

### Metabolomic age model

Untargeted NMR datasets were glog-transformed (Parsons, Ludwig, Gunther, & Viant, 2007), the quantified BiLISA data was log-transformed, and the UPLC-MS data were log transformed, following unit addition to every value to allow transformation of zero values. Data were then mean centred and scaled to unit variance.

A predictive model of metabolomic age was constructed using elastic net regression (Zou & Hastie, 2005) in the “glmnet” package (Friedman, Hastie, & Tibshirani, 2010) in R. The model was fitted on metabolic features from across all metabolomic datasets, using a multi-step process on 80% of the data (the training dataset). The remaining 20% was reserved for assessment of the predictive ability (Pearson’s correlation between predicted and chronological age) of the model in an independent dataset (the test dataset). The steps were as follows:

Step 1 Parameterisation: Elastic net model parameters, α (that defines mixing between lasso and ridge penalties) and λ (overall strength of penalty), were found following 10-fold cross validation. A line search across α, between 0 and 1 in 0.01 increments, was performed to find the minimum mean cross-validated error (MSE) using the optimal value of λ found using the ‘cvfit’ command for each α value.

Step 2 Leave platform out analysis: Due to potential redundancy between metabolomic datasets, we performed the parameterisation step above on data with one metabolomic platform left out each time. Platforms were removed from further analysis if model performed better (lower MSE) with their exclusion. We continued this process leaving further platforms out each time until no improvement in MSE was observed.

Step 3 Stability analysis: Using the selected metabolomic datasets, we repeated elastic net regression on 100 subsamples of the training dataset (a random selection of 80% each time). The metabolic features selected in each model was stored for each iteration.

Step 4 Metabolomic data restriction: On the same subsample for 101 iterations, the number of metabolic features available to build an elastic net model was restricted by the percentage of iterations in step 3 that a feature was selected, moving from 100% to 0%, in 1% decrements for each subsequent iteration. The correlation between predicted and chronological age in remaining 20% of training set was stored for each iteration and the percentage restriction value that gave the best correlation, was chosen for the final metabolic feature restriction in step 5.

Step 5 Final model building: On the complete training dataset, a final elastic net model was constructed using metabolic features restricted to those present in a set percentage of models, as found in step 4.

Metabolomic age acceleration (metAA) was defined as the difference between chronological age and predicted age, adjusted on actual age as previously defined for DNA methylation age acceleration (Horvath, 2013). That is, we define mAA as the residuals of a linear regression between the chronological age and predicted age difference, with chronological age itself.

### Metabolic feature and pathway annotation

Tentative annotations were provided for mass-spectrometry based metabolic features bases on m/z searches across the Human metabolome database (Wishart et al., 2018), for the ion forms M+2H, M+H+NH4, M+NH4, M+H, M+ACN+H, M+CH3OH+H, M+Na, M+K, 2M+H at ± 8 ppm mass tolerance.

For five UPLC-MS based metabolic features that were both tentatively annotated by exact mass within our metabolomic age model and also available in repeat measurements within the Northern Finnish Birth Cohort dataset, we performed further annotation procedures. Two of these annotations, for citrate (as in-source fragmentation product) and leucine (M+Na ionic form), were supported by matching retention times and accurate mass to an internal reference standard database.

Significantly enriched metabolic pathways were predicted using the mummichog program (Li et al., 2013). The algorithm searches tentative compound lists from metabolite reference databases against an integrated model of human metabolism to identify functional activity. Fisher’s exact tests and permutation are used to infer p-values for likelihood of pathway enrichment among significant features as compared to pathways identified among the entire compound set present in reference list (the entire metabolome dataset), considering the probability of mapping the significant m/z features to pathways. Mummichog parameters were set to match against ions included in the ‘generic positive mode’ setting at ± 8 ppm mass tolerance.

### Metabolite validation in the Northern Finnish Birth Cohort 1966

The Northern Finnish Birth Cohort 1966 is a prospective birth cohort that sampled 12,058 live births in 1966, including 96.3% of all births in the regions of Oulu and Lapland in Finland (Rantakallio, 1988). Fasting blood samples were collected at follow-up of participants at ages 31 and 46 yrs and stored at -80 °C for subsequent biomarker profiling. A high-throughput NMR metabolomics platform was used for the analysis of 87 metabolic measures (Soininen, Kangas, Wurtz, Suna, & Ala-Korpela, 2015). This metabolomics platform provides simultaneous quantification of routine lipids and lipid concentrations of 14 lipoprotein subclasses and major sub-fractions, and further quantifies abundant fatty acids, amino acids, ketone bodies and gluconeogenesis-related metabolites in absolute concentration units.

We assessed changes of nine metabolites, that were available in this dataset and also included in our predictive model, between these two sampling points using 1-tailed t-tests.

### DNA methylation analysis

For the microarray, bisulphite conversion of 500 ng of each DNA sample was performed using the EZ DNA Methylation-Lightning™ Kit according to the manufacturer’s protocol (Zymo Research, Orange, CA). Then, bisulfite-converted DNA was used for hybridization on the Infinium HumanMethylation EPIC BeadChip, following the Illumina Infinium HD Methylation protocol. Briefly, a whole genome amplification step was followed by enzymatic end-point fragmentation and hybridization to HumanMethylation EPIC BeadChips at 48°C for 17 h, followed by single nucleotide extension. The incorporated nucleotides were labelled with biotin (ddCTP and ddGTP) and 2,4-dinitrophenol (DNP) (ddATP and ddTTP). After the extension step and staining, the BeadChip was washed and scanned using the Illumina HiScan SQ scanner. The intensities of the images were extracted using the GenomeStudio (v.2011.1) Methylation module (1.9.0) software, which normalizes within-sample data using different internal controls that are present on the HumanMethylation EPIC BeadChip and internal background probes. The methylation score for each CpG was represented as a β-value according to the fluorescent intensity ratio representing any value between 0 (unmethylated) and 1 (completely methylated).

DNA methylation (DNAm) data were pre-processed and normalized using in-house software written for the R statistical computing environment, including background and color bias correction, quantile normalization, and Beta MIxture Quantile dilation (BMIQ) procedure to remove type I/type II probes bias, as described elsewhere (Fiorito et al., 2017). DNAm levels were expressed as the ratio of the intensities of methylated cytosines over the total intensities (β values). Cross-reactive and polymorphic probes - with minor allele frequency greater than 0.01 in Europeans (Y. A. Chen et al., 2013) - were excluded. Methylation measures were set to missing if the detection p-value was greater than 0.01. Samples with the bisulfite conversion control fluorescence intensity lower than 10,000 for both type I and type II probes and those with total call rate lower than 95% were excluded. Finally, samples were excluded if the predicted sex (based on chromosome X methylation) did not match that self-reported.

DNA methylation age was computed according to the algorithm described by Hannum et al. (Hannum et al., 2013) based on a set of 71 blood-specific age-associated CpG sites. We used this algorithm, rather than the algorithm of Hovarth, since it was developed specifically for blood samples and found to be the most predictive of mortality (B. H. Chen et al., 2016). Age acceleration (AA) was defined as the difference between epigenetic and chronological age. Since AA could be correlated with chronological age and WBC percentage, we computed the so-called intrinsic epigenetic age acceleration (B. H. Chen et al., 2016), which is defined as the residuals from the linear regression of AA with chronological age and blood cell counts (measured using flow cytometry) for neutrophils, lymphocytes, monocytes and eosinophils.

### Genotyping

Genotyping was performed on the Illumina Infinium HumanCoreExome-12v1-1 BeadChip and quality control filters including call rate (>=97%), heterozygosity rate (<=3SD from the mean) were applied on the samples. Duplicated and second-degree relatives were further excluded and 14,062 samples of European ancestry based on principle component analysis remained. Markers were removed for high missing rate (>2%), deviation from Hardy-Weinberg equilibrium (P<1E-5) or minor allele frequency below 1%, resulting in 254,027 high-quality and common markers. Imputation was performed using the Haplotype Reference Consortium (HRC) panel (version r1.1 2016).

We selected 10 SNPs, previously associated with parental attained age (Joshi et al., 2017; Pilling et al., 2017) from (Pilling et al., 2017), and tested their associations with both DNAmAA and mAA, in bivariate linear models. DNAmAA or mAA was used as the dependent variable and the dosage of the effect allele for each SNP (i.e. 0,1 or 2) was used as the independent variable. We also defined a weighted continuous genetic risk score calculated as described in (Pilling et al., 2017) for these 10 SNPs and tested its associations with DNAmAA and mAA in bivariate linear models.

### Analysis of risk factors of biological age acceleration

We analysed associations between mortality risk factors and age acceleration scores in separate adjusted linear regression models. To allow comparison across multiple risk factors, the adjustment set, included in all models, was chosen *a priori*. It included demographic variables (sex, ethnicity, study centre, income), the 25 x 25 main NCD risk factors, (hypertension, diabetes, BMI, smoking, alcohol intake, physical activity), and dietary indicators (DASH score and fish consumption, chosen following significant bivariate associations with mAA). Considering the exploratory nature of the analysis, we considered *p* values below 0.05 as “nominally significant” and *p* values below 0.0025 as significant after correction for multiple testing (Bonferroni-corrected for 10 risk factors x 2 outcomes).

## Acknowledgements

OR was supported by a MRC Early Career Fellowship. This study was partly supported by the European Commission grant to the LIFEPATH project (Horizon 2020 grant number 633666). The Airwave Health Monitoring Study is funded by the Home Office (grant number 780- TETRA) with additional support from the National Institute for Health Research (NIHR) Biomedical Research Centre. The Airwave Study uses the computing resources of the UK MEDical BIOinformatics partnership (UK MED-BIO supported by the Medical Research Council (MR/L01632X/1). We thank all Airwave participants for their contributions. We thank the late professor Paula Rantakallio (launch of NFBC1966), the participants in the 31 yrs and 46 yrs studies and the NFBC project center. NFBC1966 received financial support from University of Oulu Grants no. 24000692, Oulu University Hospital Grant no. 24301140, ERDF European Regional Development Fund Grant no. 539/2010 A31592, University of Oulu Grant no. 65354, Oulu University Hospital Grant no. 2/97, 8/97, Ministry of Health and Social Affairs Grant no. 23/251/97, 160/97, 190/97, National Institute for Health and Welfare, Helsinki Grant no. 54121, Regional Institute of Occupational Health, Oulu, Finland Grant no. 50621, 54231. I.K. acknowledges support from the EU PhenoMeNal project (Horizon 2020, 654241).

## Supplementary figures and tables

**Table s1:** Summary of metabolomic platforms used in analysis.

| Platform | Sample | Abbreviation | Details | N features/<br>spectral<br>resolution |
| --- | --- | --- | --- | --- |
| NMR<br>NOESY | urine | uNMR | highly robust, repeatable and precise platform for the detection of small molecules in human biofluids | 24493 |
| MS HPOS | urine | uHPOS | Hydrophilic interaction chromatography (HILIC), provides enhanced separation of small, highly polar molecules ionised in positive mode | 7325 |
| MS RPOS | urine | uRPOS | Reversed-phase chromatography targets small moderately polar molecules, ionised in positive and negative modes | 14300 |
| MS RNEG | urine | uRNEG |  | 14481 |
| NMR<br>CPMG | serum | sNMR | highly robust, repeatable and precise platform for the detection of small molecules in human biofluids | 23571 |
| NMR<br>BiLISA | serum | sBiLISA | Quantifies cholesterol, free cholesterol, phospholipids, triglycerides, apolipoproteins A1, A2, B and particle numbers for the primary plasma and serum lipoproteins and their subclasses | 105 |
| MS HPOS | serum | sHPOS | Hydrophilic interaction chromatography (HILIC), provides enhanced separation of small, highly polar molecules ionised in positive mode | 1505 |
| MS LPOS | serum | sLPOS | lipid-targeted reverse-phase chromatography provides maximal resolution of fattyacids, triglycerides, and phospholipids, , ionised in positive and negative modes | 7211 |
| MS LNEG | serum | sLNEG |  | 5833 |

**Table s2: Model predictors with coefficients**

*Please see separate excel file*.

**Table s3 Tentative annotations of model predictors**

*Please see separate excel file*.

**Table s4:**
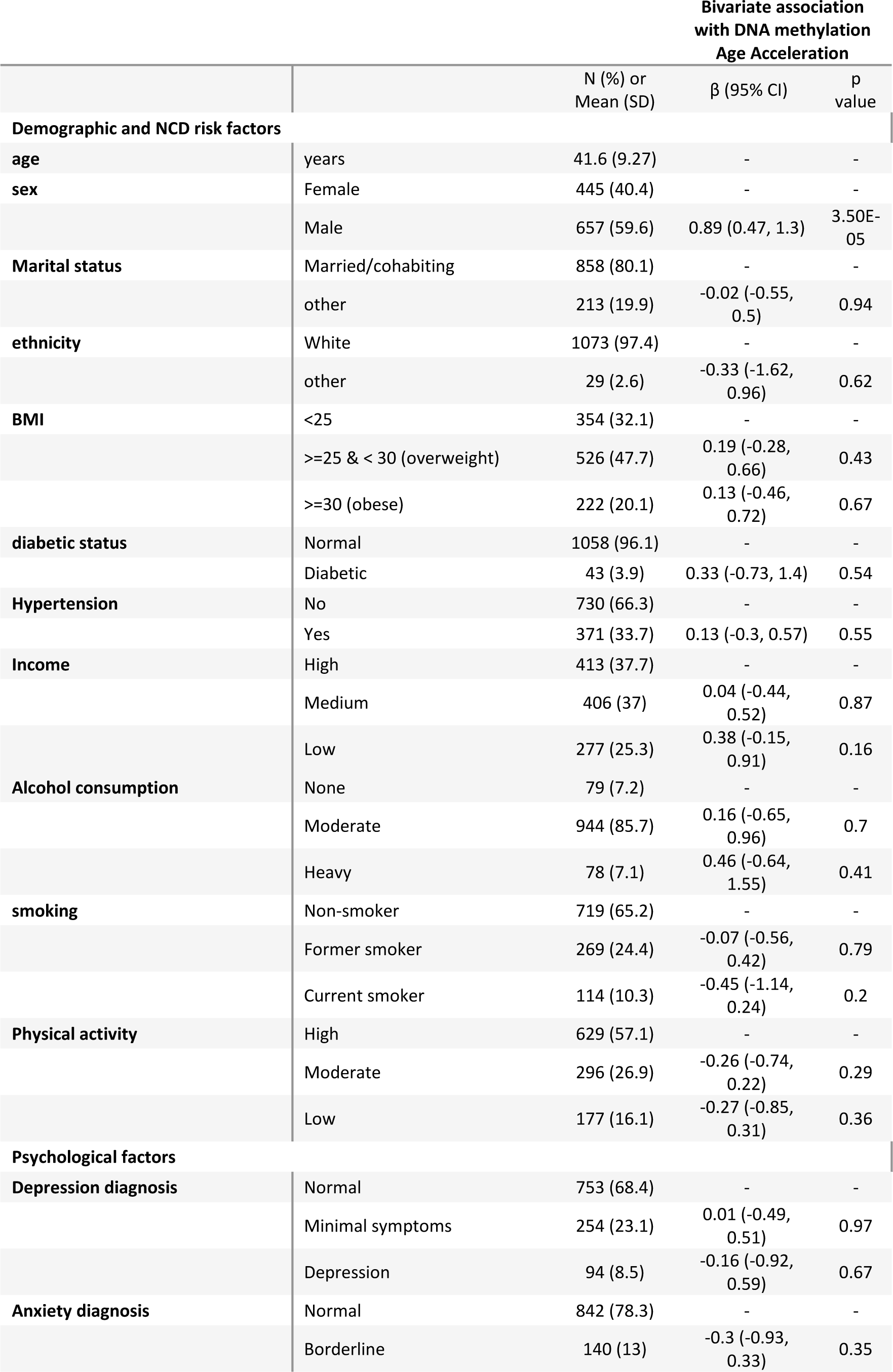
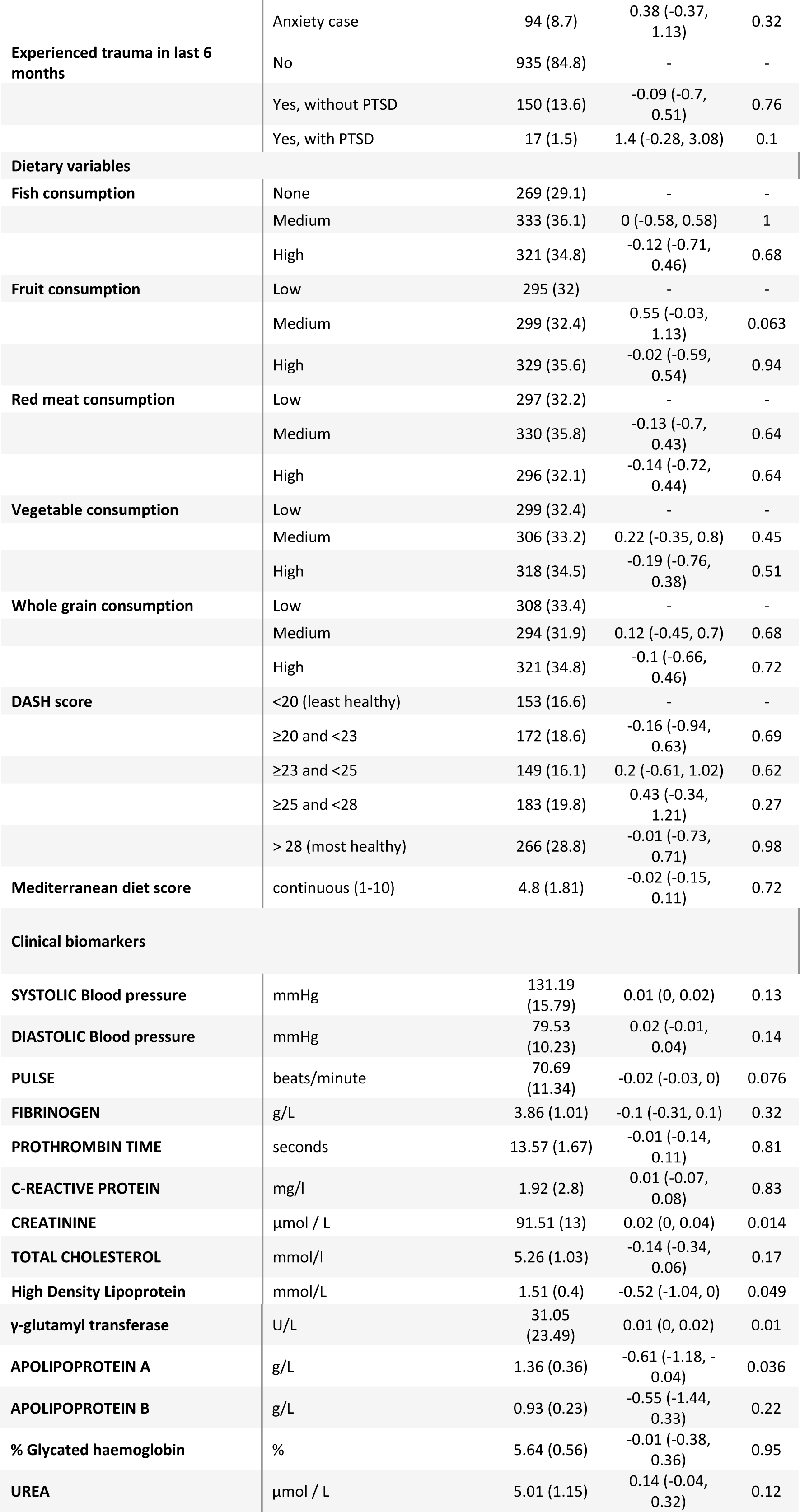
Demographic and covariate information of participants with DNA methylation data, and bivariate associations with DNA methylation age acceleration.

**Table s5:** Association of lifespan-related SNPs with age acceleration scores.

| Implicated gene(s) | SNP | CHR | BP | A1 | A0 | Association with parental lifespan<sup>a</sup> |  | Association with mAA |  | Association with DNAmAA |  |
| --- | --- | --- | --- | --- | --- | --- | --- | --- | --- | --- | --- |
|  |  |  |  |  |  | Beta | p-value | Beta (95% CI) | p-value | Beta (95% CI) | p-value |
| *CLESR2 ... PSRC1* | rs602633 | 1 | 109821511 | T | G | -0.0150 | 2.7E-08 | 0.00 (-0.26, 0.25) | 0.98 | 0.12 (-0.24, 0.48) | 0.51 |
| *HLA-DRB1... HLA-DQA1* | rs28383322 | 6 | 32592796 | C | T | 0.018 | 5.3E-11 | -0.3 (-0.57, -0.02) | 0.03 | -0.11 (-0.47, 0.26) | 0.57 |
| *LPA* | rs55730499 | 6 | 161005610 | C | T | -0.0361 | 1.7E-18 | 0.01 (-0.39, 0.4) | 0.98 | 0.14 (-0.41, 0.69) | 0.62 |
| *EPHX2* | rs7844965 | 8 | 27442064 | G | A | 0.015 | 7.7E-09 | 0.01 (-0.23, 0.26) | 0.91 | 0.05 (-0.29, 0.39) | 0.78 |
| *CDKN2B-AS1 (ANRIL)* | rs1556516 | 9 | 22100176 | G | C | -0.0181 | 4.7E-16 | 0.01 (-0.21, 0.22) | 0.95 | -0.12 (-0.41, 0.18) | 0.45 |
| *SH2B3/ATXN2* | rs7137828 | 12 | 111932800 | C | T | 0.017 | 3.4E-14 | 0.03 (-0.18, 0.24) | 0.80 | -0.02 (-0.33, 0.28) | 0.89 |
| *PROX2* | rs61978928 | 14 | 75321714 | T | C | 0.014 | 2.0E-08 | 0.02 (-0.21, 0.25) | 0.86 | 0.07 (-0.25, 0.39) | 0.66 |
| *CHRNA3* | rs1317286 | 15 | 78896129 | A | G | -0.0254 | 1.2E-26 | -0.04 (-0.27, 0.18) | 0.70 | -0.18 (-0.49, 0.13) | 0.25 |
| *FURIN* | rs17514846 | 15 | 91416550 | C | A | -0.0139 | 7.1E-10 | 0.03 (-0.18, 0.24) | 0.77 | 0.14 (-0.15, 0.43) | 0.34 |
| *APOE/APOC1* | rs429358 | 19 | 45411941 | T | C | -0.0566 | 1.4E-74 | -0.28 (-0.58, 0.01) | 0.05 | -0.38 (-0.78, 0.03) | 0.07 |

<sup>a</sup> Reproduced from (Pilling et al., 2017). For parental attained age (Martingale residuals) a negative BETA = reduced hazard, i.e. increased attained age. Implicated gene(s) = Variants in locus usually intronic, or exonic (indicated by underlining). If two genes separated by dots, the SNP is intergenic. BP = Genomic position, build 37 (hg19). Betas = A1 effect on outcome.

**Figure s1:**
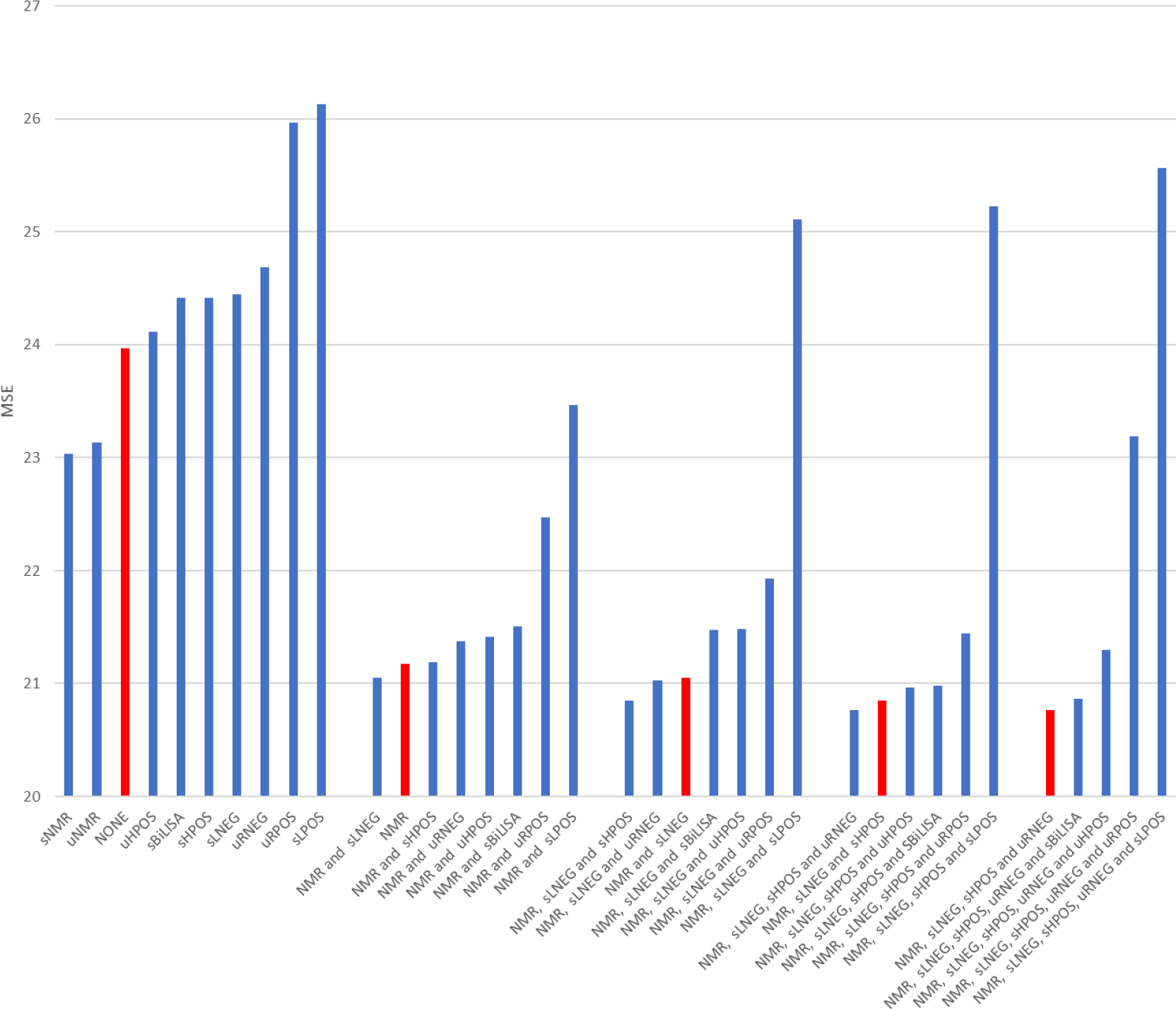
Results of &leave platform out’ stage of model building

## References

Auro, K., Joensuu, A., Fischer, K., Kettunen, J., Salo, P., Mattsson, H.,… Perola, M. (2014). A metabolic view on menopause and ageing. Nature Communications, 5, 1–11. doi:10.1038/ncomms5708

Belsky, D. W., Caspi, A., Houts, R., Cohen, H. J., Corcoran, D. L., Danese, A.,… Moffitt, T. E. (2015). Quantification of biological aging in young adults. Proc.Natl.Acad.Sci.U.S.A, 112(30), E4104– E4110. doi:10.1073/pnas.1506264112

Belsky, D. W., Moffitt, T. E., Cohen, A. A., Corcoran, D. L., Horvath, S., Levine, M. E.,… Caspi, A. (2016). Telomere, epigenetic clock, and biomarker-composite quantifications of biological aging: Do they measure the same thing? bioRxiv, 071373-071373. doi:10.1101/071373

Brewin, C. R., Rose, S., Andrews, B., Green, J., Tata, P., McEvedy, C.,… Foa, E. B. (2002). Brief screening instrument for post-traumatic stress disorder. Br J Psychiatry, 181, 158–162.

Brown, P. J., Wall, M. M., Chen, C., Levine, M. E., Yaffe, K., Roose, S. P., & Rutherford, B. R. (2017). Biological Age, Not Chronological Age, is Associated with Late Life Depression. The Journals of Gerontology: Series A, 00(00), 1–7. doi:10.1093/gerona/glx162

Burkle, A., Moreno-Villanueva, M. a., Bernhard, J. r., Blasco, M. a., Zondag, G., Hoeijmakers, J. H. J.,… Aspinall, R. (2015). MARK-AGE biomarkers of ageing. Mechanisms of Ageing and Development, 151, 2–12. doi:10.1016/j.mad.2015.03.006

Castagne, R., Gares, V., Karimi, M., Chadeau-Hyam, M., Vineis, P., Delpierre, C., & Kelly-Irving, M. (2018). Allostatic load and subsequent all-cause mortality: which biological markers drive the relationship? Findings from a UK birth cohort. Eur J Epidemiol. doi:10.1007/s10654-018-0364-1

Chaleckis, R., Murakami, I., Takada, J., Kondoh, H., & Yanagida, M. (2016). Individual variability in human blood metabolites identifies age-related differences. Proceedings of the National Academy of Sciences of the United States of America, 113(16), 4252–4259. doi:10.1073/pnas.1603023113

Chang, H.-C., & Guarente, L. (2013). SIRT1 Mediates Central Circadian Control in the SCN by a Mechanism that Decays with Aging. Cell, 153(7), 1448–1460. doi:https://doi.org/10.1016/j.cell.2013.05.027

Chen, B. H., Marioni, R. E., Colicino, E., Peters, M. J., Ward-Caviness, C. K., Tsai, P. C.,… Horvath, S. (2016). DNA methylation-based measures of biological age: meta-analysis predicting time to death. Aging (Albany NY), 8(9), 1844–1865. doi:10.18632/aging.101020

Chen, Y. A., Lemire, M., Choufani, S., Butcher, D. T., Grafodatskaya, D., Zanke, B. W.,… Weksberg, R. (2013). Discovery of cross-reactive probes and polymorphic CpGs in the Illumina Infinium HumanMethylation450 microarray. Epigenetics, 8(2), 203–209. doi:10.4161/epi.23470

Chiu, M., Vigod, S., Rahman, F., Wilton, A. S., Lebenbaum, M., & Kurdyak, P. (2018). Mortality risk associated with psychological distress and major depression: A population-based cohort study. Journal of Affective Disorders, 234(February), 117-123. doi:10.1016/j.jad.2018.02.075

Claycombe, K. J., & Meydani, S. N. (2001). Vitamin E and genome stability. Mutat Res, 475(1-2), 37–44.

De Jager, P. L., Shulman, J. M., Chibnik, L. B., Keenan, B. T., Raj, T., Wilson, R. S.,… Evans, D. A. (2012). A genome-wide scan for common variants affecting the rate of age-related cognitive decline. Neurobiol Aging, 33(5), 1017.e1011-1015. doi:10.1016/j.neurobiolaging.2011.09.033

Dona, A. C., Jimenez, B., Schafer, H., Humpfer, E., Spraul, M., Lewis, M. R.,… Nicholson, J. K. (2014). Precision high-throughput proton NMR spectroscopy of human urine, serum, and plasma for large-scale metabolic phenotyping. Anal Chem, 86(19), 9887–9894. doi:10.1021/ac5025039

Dowd, J. B., Simanek, A. M., & Aiello, A. E. (2009). Socio-economic status, cortisol and allostatic load: a review of the literature. Int J Epidemiol, 38(5), 1297–1309. doi:10.1093/ije/dyp277

Dugue, P. A., Bassett, J. K., Joo, J. E., Baglietto, L., Jung, C. H., Wong, E. M.,… Milne, R. L. (2018). Association of DNA Methylation-Based Biological Age With Health Risk Factors and Overall and Cause-Specific Mortality. Am J Epidemiol, 187(3), 529–538. doi:10.1093/aje/kwx291

Elliott, P., Vergnaud, A.-C., Singh, D., Neasham, D., Spear, J., & Heard, A. (2014). The Airwave Health Monitoring Study of police officers and staff in Great Britain: Rationale, design and methods. Environmental research, 134C, 280-285. doi:10.1016/j.envres.2014.07.025

Fang, Evandro F., Scheibye-Knudsen, M., Brace, Lear E., Kassahun, H., SenGupta, T., Nilsen, H.,… Bohr, Vilhelm A. (2014). Defective Mitophagy in XPA via PARP-1 Hyperactivation and NAD+/SIRT1 Reduction. Cell, 157(4), 882–896. doi:https://doi.org/10.1016/j.cell.2014.03.026

Fiorito, G., Polidoro, S., Dugue, P. A., Kivimaki, M., Ponzi, E., Matullo, G.,… Vineis, P. (2017). Social adversity and epigenetic aging: a multi-cohort study on socioeconomic differences in peripheral blood DNA methylation. Sci Rep, 7(1), 16266. doi:10.1038/s41598-017-16391-5

Friedman, J., Hastie, T., & Tibshirani, R. (2010). Regularization Paths for Generalized Linear Models via Coordinate Descent. J Stat Softw, 33(1), 1–22.

Fung, T. T., Chiuve, S. E., McCullough, M. L., Rexrode, K. M., Logroscino, G., & Hu, F. B. (2008). Adherence to a DASH-style diet and risk of coronary heart disease and stroke in women. Arch Intern Med, 168(7), 713–720. doi:10.1001/archinte.168.7.713

Garinis, G. A., van der Horst, G. T. J., Vijg, J., & H.J. Hoeijmakers, J. (2008). DNA damage and ageing: new-age ideas for an age-old problem. Nature Cell Biology, 10, 1241. doi:10.1038/ncb1108-1241

Gibson, R., Eriksen, R., Lamb, K., McMeel, Y., Vergnaud, A. C., Spear, J.,… Frost, G. (2017). Dietary assessment of British police force employees: a description of diet record coding procedures and cross-sectional evaluation of dietary energy intake reporting (The Airwave Health Monitoring Study). BMJ Open, 7(4), e012927. doi:10.1136/bmjopen-2016-012927

Gomes, Ana P., Price, Nathan L., Ling, Alvin J. Y., Moslehi, Javid J., Montgomery, M. K., Rajman, L.,… Sinclair, David A. (2013). Declining NAD+ Induces a Pseudohypoxic State Disrupting Nuclear-Mitochondrial Communication during Aging. Cell, 155(7), 1624–1638. doi:https://doi.org/10.1016/j.cell.2013.11.037

Hannum, G., Guinney, J., Zhao, L., Zhang, L., Hughes, G., Sadda, S.,… Zhang, K. (2013). Genome-wide methylation profiles reveal quantitative views of human aging rates. Mol Cell, 49(2), 359–367. doi:10.1016/j.molcel.2012.10.016

Hertel, J., Friedrich, N., Wittfeld, K., Pietzner, M., Budde, K., Van Der Auwera, S.,… Grabe, H. J. (2016). Measuring Biological Age via Metabonomics: The Metabolic Age Score. Journal of Proteome Research, 15(2), 400–410. doi:10.1021/acs.jproteome.5b00561

Horvath, S. (2013). DNA methylation age of human tissues and cell types. Genome Biology, 14(10), R115–R115. doi:10.1186/gb-2013-14-10-r115

Horvath, S., Gurven, M., Levine, M. E., Trumble, B. C., Kaplan, H., Allayee, H.,… Assimes, T. L. (2016). An epigenetic clock analysis of race/ethnicity, sex, and coronary heart disease. Genome Biology, 17(1), 0–22. doi:10.1186/s13059-016-1030-0

Ito, K., Hirao, A., Arai, F., Takubo, K., Matsuoka, S., Miyamoto, K.,… Suda, T. (2006). Reactive oxygen species act through p38 MAPK to limit the lifespan of hematopoietic stem cells. Nature Medicine, 12, 446. doi:10.1038/nm1388

Joshi, P. K., Pirastu, N., Kentistou, K. A., Fischer, K., Hofer, E., Schraut, K. E.,… Wilson, J. F. (2017). Genome-wide meta-analysis associates HLA-DQA1/DRB1 and LPA and lifestyle factors with human longevity. Nature Communications, 8(1), 910. doi:10.1038/s41467-017-00934-5

Jylhävä, J., Pedersen, N. L., & Hägg, S. (2017). Biological Age Predictors (Vol. 21, pp. 29-36): The Authors.

Kroenke, K., Spitzer, R. L., & Williams, J. B. (2001). The PHQ-9: validity of a brief depression severity measure. J Gen Intern Med, 16(9), 606–613.

Laimer, M., Melmer, A., Lamina, C., Raschenberger, J., Adamovski, P., Engl, J.,… Ebenbichler, C. (2015). Telomere length increase after weight loss induced by bariatric surgery: results from a 10 year prospective study. International Journal Of Obesity, 40, 773. doi:10.1038/ijo.2015.238

Levine, M. E. (2013). Modeling the rate of senescence: Can estimated biological age predict mortality more accurately than chronological age? Journals of Gerontology - Series A Biological Sciences and Medical Sciences, 68(6), 667–674. doi:10.1093/gerona/gls233

Lewis, M. R., Pearce, J. T. M., Spagou, K., Green, M., Dona, A. C., Yuen, A. H. Y.,… Nicholson, J. K. (2016). Development and Application of Ultra-Performance Liquid Chromatography-TOF MS for Precision Large Scale Urinary Metabolic Phenotyping. Analytical Chemistry, 88(18), 9004–9013. doi:10.1021/acs.analchem.6b01481

Li, S., Park, Y., Duraisingham, S., Strobel, F. H., Khan, N., Soltow, Q. A.,… Pulendran, B. (2013). Predicting Network Activity from High Throughput Metabolomics. PLOS Computational Biology, 9(7), e1003123. doi:10.1371/journal.pcbi.1003123

López-Otín, C., Blasco, M. A., Partridge, L., Serrano, M., & Kroemer, G. (2013). The hallmarks of aging (Vol. 153).

López-Otín, C., Galluzzi, L., Freije, J. M. P., Madeo, F., & Kroemer, G. (2016). Metabolic Control of Longevity (Vol. 166, pp. 802–821).

Marazziti, D., Rutigliano, G., Baroni, S., Landi, P., & Dell’Osso, L. (2014). Metabolic syndrome and major depression (Vol. 19, pp. 293–304).

Marioni, R. E., Harris, S. E., Zhang, Q., McRae, A. F., Hagenaars, S. P., Hill, W. D.,… Visscher, P. M. (2018). GWAS on family history of Alzheimer’s disease. Transl Psychiatry, 8(1), 99. doi:10.1038/s41398-018-0150-6

McDaid, A. F., Joshi, P. K., Porcu, E., Komljenovic, A., Li, H., Sorrentino, V.,… Kutalik, Z. (2017). Bayesian association scan reveals loci associated with human lifespan and linked biomarkers. Nature Communications, 8(May). doi:10.1038/ncomms15842

McEwen, B. S., & Seeman, T. (1999). Protective and damaging effects of mediators of stress. Elaborating and testing the concepts of allostasis and allostatic load. Ann N Y Acad Sci, 896, 30–47.

Moffitt, T. E., Belsky, D. W., Danese, A., Poulton, R., & Caspi, A. (2017). The Longitudinal Study of Aging in Human Young Adults: Knowledge Gaps and Research Agenda. The journals of gerontology. Series A, Biological sciences and medical sciences, 72(2), 210–215. doi:10.1093/gerona/glw191

Muezzinler, A., Zaineddin, A. K., & Brenner, H. (2013). A systematic review of leukocyte telomere length and age in adults. Ageing Res Rev, 12(2), 509–519. doi:10.1016/j.arr.2013.01.003

Parsons, H. M., Ludwig, C., Gunther, U. L., & Viant, M. R. (2007). Improved classification accuracy in 1- and 2-dimensional NMR metabolomics data using the variance stabilising generalised logarithm transformation. BMC Bioinformatics, 8, 234. doi:10.1186/1471-2105-8-234

Partridge, L., Deelen, J., & Slagboom, P. E. (2018). Facing up to the global challenges of ageing. Nature, 561(7721), 45–56. doi:10.1038/s41586-018-0457-8

Peters, M. J., Joehanes, R., Pilling, L. C., Schurmann, C., Conneely, K. N., Powell, J.,… Singleton, A. B. (2015). The transcriptional landscape of age in human peripheral blood. Nature Communications, 6. doi:10.1038/ncomms9570

Pilling, L. C., Kuo, C. L., Sicinski, K., Tamosauskaite, J., Kuchel, G. A., Harries, L. W.,… Melzer, D. (2017). Human longevity: 25 genetic loci associated in 389,166 UK biobank participants. Aging (Albany NY), 9(12), 2504–2520. doi:10.18632/aging.101334

Rantakallio, P. (1988). The longitudinal study of the northern Finland birth cohort of 1966. Paediatr Perinat Epidemiol, 2(1), 59–88.

Rist, M. J., Roth, A., Frommherz, L., Weinert, C. H., Krüger, R., Merz, B.,… Watzl, B. (2017). Metabolite patterns predicting sex and age in participants of the Karlsruhe Metabolomics and Nutrition (KarMeN) study. PLoS ONE, 12(8), 1–21. doi:10.1371/journal.pone.0183228

Sahar, S., Masubuchi, S., Eckel-Mahan, K., Vollmer, S., Galla, L., Ceglia, N.,… Sassone-Corsi, P. (2014). Circadian Control of Fatty Acid Elongation by SIRT1 Protein-mediated Deacetylation of Acetyl-coenzyme A Synthetase 1. Journal of Biological Chemistry, 289(9), 6091–6097. doi:10.1074/jbc.M113.537191

Sala, M. L., Röell, B., van der Bijl, N., van der Grond, J., de Craen, A. J. M., Slagboom, E. P.,… Kroft, L. J. M. (2015). Genetically determined prospect to become long-lived is associated with less abdominal fat and in particular less abdominal visceral fat in men. Age and Ageing, 44(4), 713–717. doi:10.1093/ageing/afv063

Sarafian, M. H., Gaudin, M., Lewis, M. R., Martin, F. P., Holmes, E., Nicholson, J. K., & Dumas, M. E. (2014). Objective set of criteria for optimization of sample preparation procedures for ultra-high throughput untargeted blood plasma lipid profiling by ultra performance liquid chromatography-mass spectrometry. Anal Chem, 86(12), 5766–5774. doi:10.1021/ac500317c

Soininen, P., Kangas, A. J., Wurtz, P., Suna, T., & Ala-Korpela, M. (2015). Quantitative serum nuclear magnetic resonance metabolomics in cardiovascular epidemiology and genetics. Circ Cardiovasc Genet, 8(1), 192–206. doi:10.1161/circgenetics.114.000216

Stringhini, S., Carmeli, C., Jokela, M., Avendano, M., Muennig, P., Guida, F.,… Kivimaki, M. (2017). Socioeconomic status and the 25 x 25 risk factors as determinants of premature mortality: a multicohort study and meta-analysis of 1.7 million men and women. Lancet, 389(10075), 1229–1237. doi:10.1016/s0140-6736(16)32380-7

Su, X., Wellen, K. E., & Rabinowitz, J. D. (2016). Metabolic control of methylation and acetylation. Current Opinion in Chemical Biology, 30, 52–60. doi:https://doi.org/10.1016/j.cbpa.2015.10.030

The IPAQ group. (2016). International Physical Activity Questionnaire. Retrieved from http://www.ipaq.ki.se/

Trichopoulou, A., Costacou, T., Bamia, C., & Trichopoulos, D. (2003). Adherence to a Mediterranean diet and survival in a Greek population. N Engl J Med, 348(26), 2599–2608. doi:10.1056/NEJMoa025039

Verdin, E. (2015). NAD+ in aging, metabolism, and neurodegeneration. Science, 350(6265), 1208– 1213. doi:10.1126/science.aac4854

Wishart, D. S., Feunang, Y. D., Marcu, A., Guo, A. C., Liang, K., Vazquez-Fresno, R.,… Scalbert, A. (2018). HMDB 4.0: the human metabolome database for 2018. Nucleic Acids Res, 46(D1), D608–d617. doi:10.1093/nar/gkx1089

Wolf, E. J., & Morrison, F. G. (2017). Traumatic Stress and Accelerated Cellular Aging: From Epigenetics to Cardiometabolic Disease (Vol. 19): Current Psychiatry Reports.

World Health Organisation. (2013). Global action plan for the prevention and control of noncommunicable diseases 2013–2020. Retrieved from

Yu, Z., Zhai, G., Singmann, P., He, Y., Xu, T., Prehn, C.,… Wang-Sattler, R. (2012). Human serum metabolic profiles are age dependent. Aging Cell, 11(6), 960–967. doi:10.1111/j.1474-9726.2012.00865.x

Zelena, E., Dunn, W. B., Broadhurst, D., Francis-McIntyre, S., Carroll, K. M., Begley, P.,… Kell, D. B. (2009). Development of a robust and repeatable UPLC-MS method for the long-term metabolomic study of human serum. Anal Chem, 81(4), 1357–1364. doi:10.1021/ac8019366

Zhao, L. G., Sun, J. W., Yang, Y., Ma, X., Wang, Y. Y., & Xiang, Y. B. (2016). Fish consumption and all-cause mortality: a meta-analysis of cohort studies. Eur J Clin Nutr, 70(2), 155–161. doi:10.1038/ejcn.2015.72

Zigmond, A. S., & Snaith, R. P. (1983). The hospital anxiety and depression scale. Acta Psychiatr Scand, 67(6), 361–370.

Zou, H., & Hastie, T. (2005). Regularization and variable selection via the elastic net. Journal of the Royal Statistical Society: Series B (Statistical Methodology), 67(2), 301–320. doi:10.1111/j.1467-9868.2005.00503.x

